# High-resolution temporal profiling of the gut microbiome through the whole life of mice

**DOI:** 10.1101/2022.11.07.515511

**Authors:** Lena Takayasu, Eiichiro Watanabe, Taichi Umeyama, Rina Kurokawa, Rie Maskawa, Sumanta Basu, Yusuke Ogata, Yuya Kiguchi, Hiroaki Masuoka, Masahiro Umezaki, Hideki Takayasu, Misako Takayasu, Masahira Hattori, Wataru Suda

## Abstract

The temporal changes of the gut microbiome are thought to be critical for understanding its interactions with host aging, but lifelong dynamics within the same individual remain largely unknown. Here we firstly report the high temporal resolution dynamics of gut microbiomes in mice sharing the same genetic background and environment from their birth to natural death, spanning >1,000 days. The 16S rRNA sequencing analysis revealed 9 patterns of OTU temporal dynamics and 38 common “life-core” bacterial species/operational taxonomic units (OTUs) in ≥80% of all samples across the lifespan of individual mice. The life-core OTUs are largely represented by the phylum Bacteroidota, whereas the transient bacterial group predominantly includes the phylum Firmicutes (Bacillota). Despite the shared genetic background and dietary habits, the gut microbiome structure significantly diversified with age and among individuals. A positive correlation existed between longevity and the microbiome α-diversity in middle age (200–500 days) followed by a negative correlation in old age (>700 days), likely influenced by the increase in diversity during the last days of life. The abundance of several “life-core” species also exhibited non-static correlation trends with lifespan. Overall, this research characterized the gut microbiomes based on its persistence over host’s lifetime and suggested a non-static host-microbiome relationship within individual mice.

## Introduction

Throughout the life from birth to death, interactions between gut microbiomes and their hosts are continuous and dynamic, shaped by various factors such as diet, lifestyle and the aging process(1–4). Recent research has highlighted the importance of this lifelong symbiosis between hosts and their gut microbiota, which underlies behind health and disease across various stages of life(5–12). However, in cohort-based studies, especially in humans, it is often challenging to distinguish the effects of the natural aging process from the various environmental and social factors that each cohort has experienced uniquely, such as medical advancements, shifts in dietary habits, socioeconomic changes. There are several studies that follow a single cohort over time(13–18), leading to a more precise understanding of how the microbiome evolves within individuals, reducing the influence of generational or lifestyle differences that vary across cohorts. Despite these advancements, few studies have comprehensively tracked an individual’s microbiome from birth to natural death comprehensively(19,20). A full lifespan study would provide a unified powerful window into how gut microbiomes influence our biology throughout life.

In this study, we applied high-resolution time-course observations to evaluate the basic dynamics and stability of the gut microbiome throughout the lifespan of mice. To this end, we analyzed fecal samples collected longitudinally from siblings of specific pathogen-free (SPF) mice throughout their lives, from birth to death, spanning over 1,000 days. We described the typical dynamics and persistence of OTUs, and found the association of gut microbiomes with the lifespan of hosts and life events, including pregnancy, delivery, and cohousing, under the same rearing environments. Although the sample size was limited, our data suggest that the relationship between the host and the gut microbiome is non-static; rather, it changes across different stages of life within the same individual.

## Results

### Mice, fecal sample collection, and 16S rRNA gene analysis

We purchased and bred the parent mice (Mo and Fa, SPF C57BL/6J strain), which thereafter had nonuplet mice (one male, seven females, and one unknown) at the animal facility (RIKEN, Kanagawa, Japan). Of these siblings, seven mice (M1, M2, M3, M4, M5, M7, and M8) died of natural causes with a mean lifetime of 922.7 ± 71.6 days (range: 827–1044 days); one mouse (M6) died at 524 days during the sampling process, likely due to the debility associated with tumor growth, and was removed from the lifespan-related analysis. Additionally, one mouse (M9) died immediately after birth and was, thus, removed from analysis (**Fig. 1a**).

**Fig. 1.**
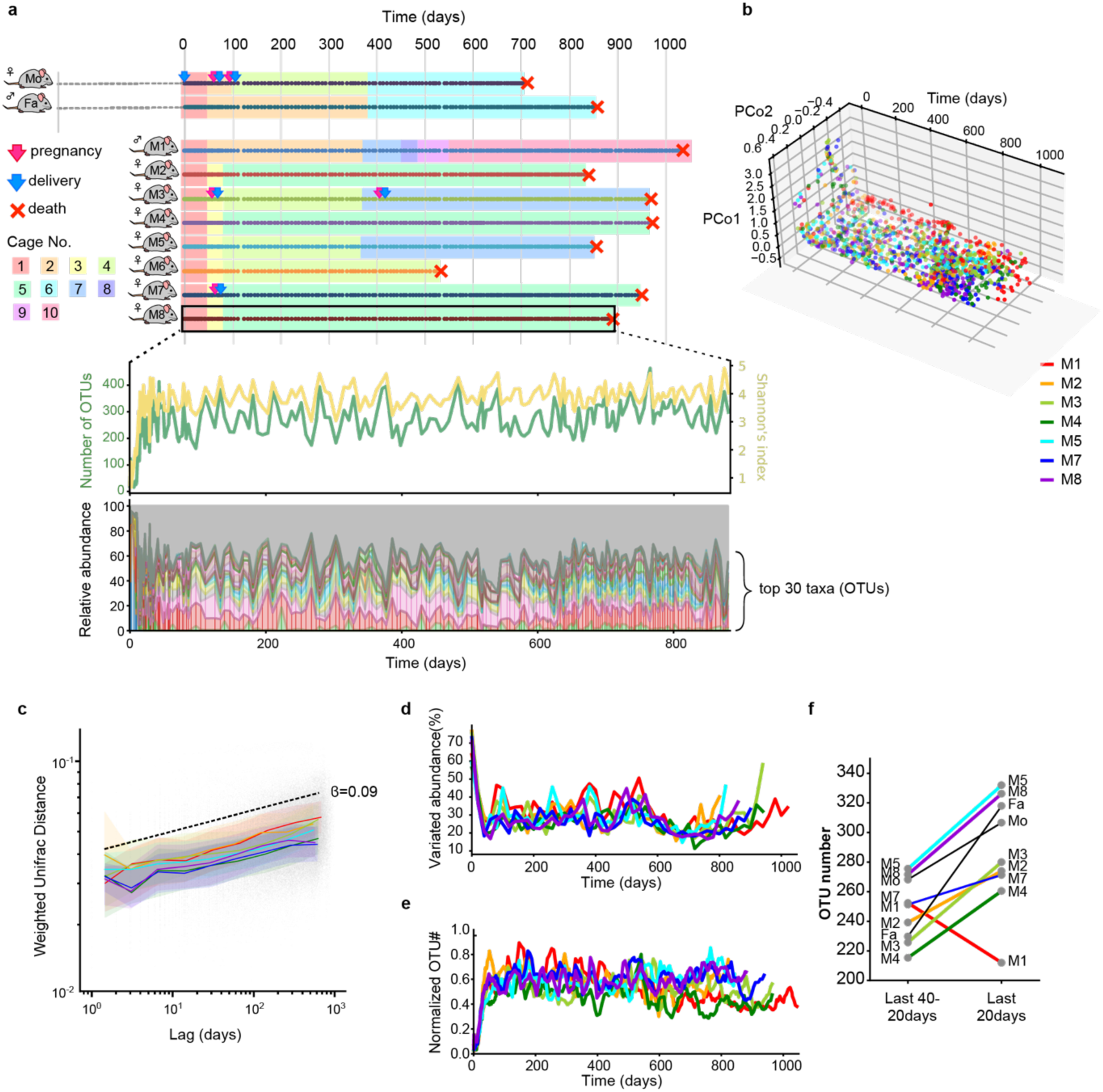
Overview of the experimental design and gut microbial diversities. **a,** Overview image of the experimental design. We performed 16S analysis on fecal samples obtained from detailed sampling, calculating diversity and the abundance of each OTU. The colored dots in the upper figure indicate the sampling points. **b,** PCoA of seven mice from day 0 to day 1044 of life based on the weighted UniFrac distance. Color corresponds to the mice. **c,** Weighted Unifrac distance within individual samples, plotted according to the time lag between samples from day 50 to day 1044. Median distances were shown in lines, and quartiles were shown in bands. Color corresponds to the mice. **d,** Temporal variations in microbiomes within individuals (See methods) with weighted Jaccard distance indicating the average change in relative abundance. Color corresponds to the mice. **e,** Simple moving average (20-days window) for OTU number throughout life, normalized by maximum OTU number for each mouse. **f,** Average OTU number of the last 40–20 days and last 20 days of life for each mouse.

Throughout the experiment, the eight siblings were cohoused with their parents for 51 days after birth and thereafter cohoused with different combinations of individuals in different cages. Additionally,three offspring mice (M3, M7, and M8) and Mo experienced pregnancy and delivery (**Fig. 1a**). Immediately upon death, we performed autopsies of the mice. Tumors were detected in five mice (M2, M6, M7, M8, and Mo).

We longitudinally collected a total of 1,815 fecal samples from ten mice (eight siblings and parents) with an average sampling interval of 4.3 days. From the fecal DNA samples, 18.0 million high-quality reads of the 16S rRNA gene V1–2 region were obtained using the Illumina MiSeq platform (Supplementary Table 1, Methods). Clustering of the 16S reads from all samples of the ten mice (eight siblings and parents) generated a total of 21,768 OTUs, including 6,531 with ≥0.1% abundance in at least one sample.

### Temporal variations in β-diversity across a lifetime

Principal coordinate analysis (PCoA) of weighted UniFrac distances of all samples across the lifetime of the seven siblings that died of natural causes revealed the most prominent difference in β-diversity during the first 50 days (**Fig. 1b**). After day 50, the UniFrac distance was observed to increase with a power law of β = 0.09 over the time lag, indicating that the changes in the microbial community followed a consistent pattern (**Fig. 1c**). Meanwhile, temporal variations in microbiomes within individuals, assessed every 20 days, revealed that the speed of microbial community changes stabilized at a relatively constant pace up to around day 500, following an initial rapid shift shortly after birth. Between days 600 and 800, the changes slowed slightly, after which an increase was observed again near the end of life (**Fig. 1d**). Permutational multivariate analysis of variance (PEAMANOVA) further revealed that the greatest dissociation in gut microbiomes was associated with time, followed by inter-individual and cage differences in unweighted UniFrac distances (seven siblings and M6; **Fig. S1a**). Meanwhile, weighted UniFrac distances (seven siblings and M6; **Fig. S1b**) showed the greatest dissociation by individuals after 21 days, followed by time and cage differences. F-values of the cage effect showed the least dissociation in weighted and unweighted UniFrac distances; however, the P-values were significantly low (*P* = 0.001).

### Temporal variations in α-diversity across a lifetime

The number of OTUs rapidly increased the first 20 days, then showed a gradual decrease or relatively constant after 100 days in most mice (**Figs. 1e and S2a**). Analysis of the cumulative number of major OTUs (≥0.1% abundance in at least one sample) over a 200-day lifetime from birth revealed that over 50% and 90% of the total OTUs appeared within 28 and 167 days in the samples of the seven siblings, respectively (**Fig. S2b**). We also observed an increase in the average number of OTUs within the final 20 days before death in eight out of nine mice, including parent mice (**Fig. 1f**).

### Variations in the relative abundance of OTUs across a lifetime

We classified the abundant OTUs (0.1% abundance across lifetimes for each mouse) in the samples from seven siblings into 9 clusters based on hierarchical clustering of temporal dynamics (**Figs. 2a, S3– S4**). The number of OTUs in each cluster, from Cluster_1 to Cluster_9, was 45, 100, 100, 25, 58, 23, 58, 213, and 84 (counting common OTUs different mice), respectively. Cluster_1, 4 and 5 commonly showed transient profiles in the first 50 days. Cluster_2 included a larger portion of Bacteroidota OTUs and exhibited highest average persistency (detected frequency) among all clusters. Cluster_6, 7 and 9 contained OTUs that were more abundant in the first half of life, while Cluster_3 and 8 comprised OTUs that were relatively abundant in the latter half. The major OTUs (>5% maximum abundance) that appeared in the first 20 days are listed in **Fig. S5**; Cluster_4 was primarily composed of *Enterococcaceae,* and *Streptococcaceae* within the first 6 days, followed by other Lactobacillaceae from day 8 and Bacteroidota species from day 14.

**Fig. 2.**
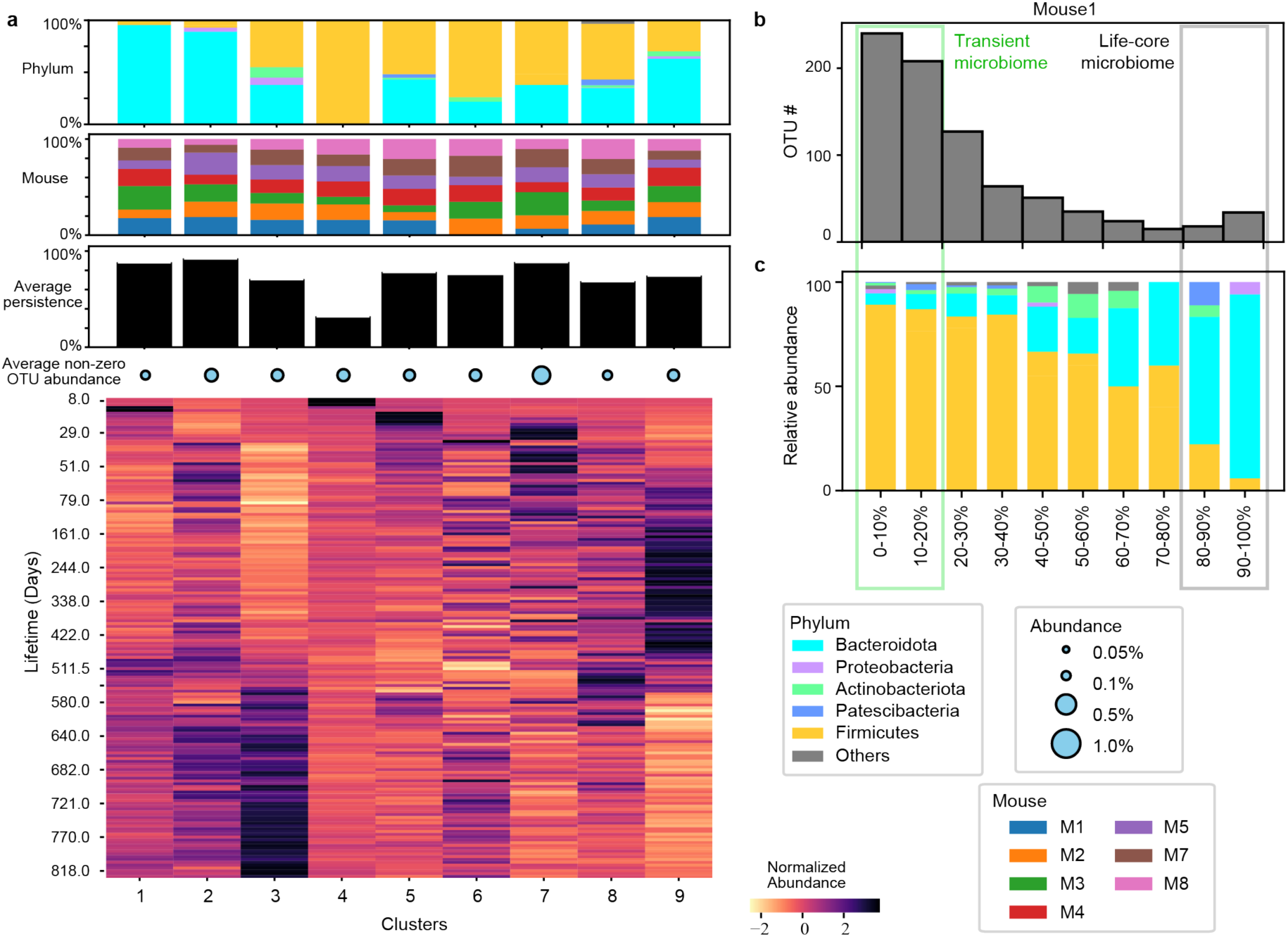
Life-core and transient microbiome. **a,** 9 clusters of temporal OTU dynamics (see Methods). Stacked bar graphs indicating the phylum-level composition (top), the ratio of OTUs from each mouse (middle), and average persistence of OTUs included in the cluster (bottom). The heatmap shows the average relative abundance of OTUs within each temporal dynamic cluster. **b,** Histograms of the observed frequency of the OTUs (maximum relative abundance ≥ 0.1%). Average persistence was calculated as the proportion of observations with abundance >0 reads relative to the total number of observations. **c,** Stacked bar graphs indicating the phylum-level composition according to the persistence.

We also identified the change-point for each OTU as the time point that caused the largest shift in the mean abundance before and after. While many OTUs exhibited change-points within the first 100 days, a period between 300 and 800 days was also observed as a phase during which non-stationary changes frequently occurred (**Fig. S6**).

Histograms depicting the number of OTUs (≥0.1% abundance in at least one sample) according to their persistence throughout the life showed a bimodal distribution in all samples of the seven siblings (**Figs. 2b, S7**). We then defined OTUs detected in >80% and <20% of all samples from each mouse as “life-core” and “transient” OTUs, respectively.

We found 69 non-redundant life-core OTUs (ranging from 48 to 60 across the seven siblings), of which 38 (minimum life-core OTUs, 51%) were common in all mice, and 4 OTUs were specific to individuals (Supplementary Table 2). Moreover, the proportion of the life-core OTUs was highest in Clusters_2 (93.0%). In contrast, 989 non-redundant transient OTUs (ranging from 317 to 448 across the seven siblings), of which 394/989 (39.8%) were specific to individual mice.

Taxonomic assignment revealed that the most of the life-core OTUs assigned to Bacteroidota in the lifecore, while the transient OTUs were predominantly assigned to Firmicutes (Bacillota) (**Fig. 2c, Fig. S7**). In addition, life-core OTUs had a relatively high abundance compared to transient OTUs (**Fig. S8**). The total abundances of life-core and transient OTUs through life are shown in **Fig. S9**. In early life, transient OTUs were highly abundant, whereas their abundance decreased to <10 % in late life. Meanwhile, the life-core OTUs dominated 50%–80% of the total abundance throughout the majority of the murine lifespan.

### Nonstationary correlation with host lifespan

The seven siblings had varying lifespans ranging from 832 to 1,049 days (Supplementary Table 1), prompting us to explore the correlation between gut microbiome structure and lifespan.

Considering the long-term transition of the gut microbiome structure, we assessed the correlation between lifespan and average α-diversities at various time points and duration (**Fig. S10**). The correlation of average diversities throughout the life was not strongly correlated with lifespan (top right corner, **Fig. S10**). However, positive correlations were detected in middle age (days 200–500), whereas a negative correlation was observed in old age (after day 700) (**Fig. 3a**).

**Fig.3.**
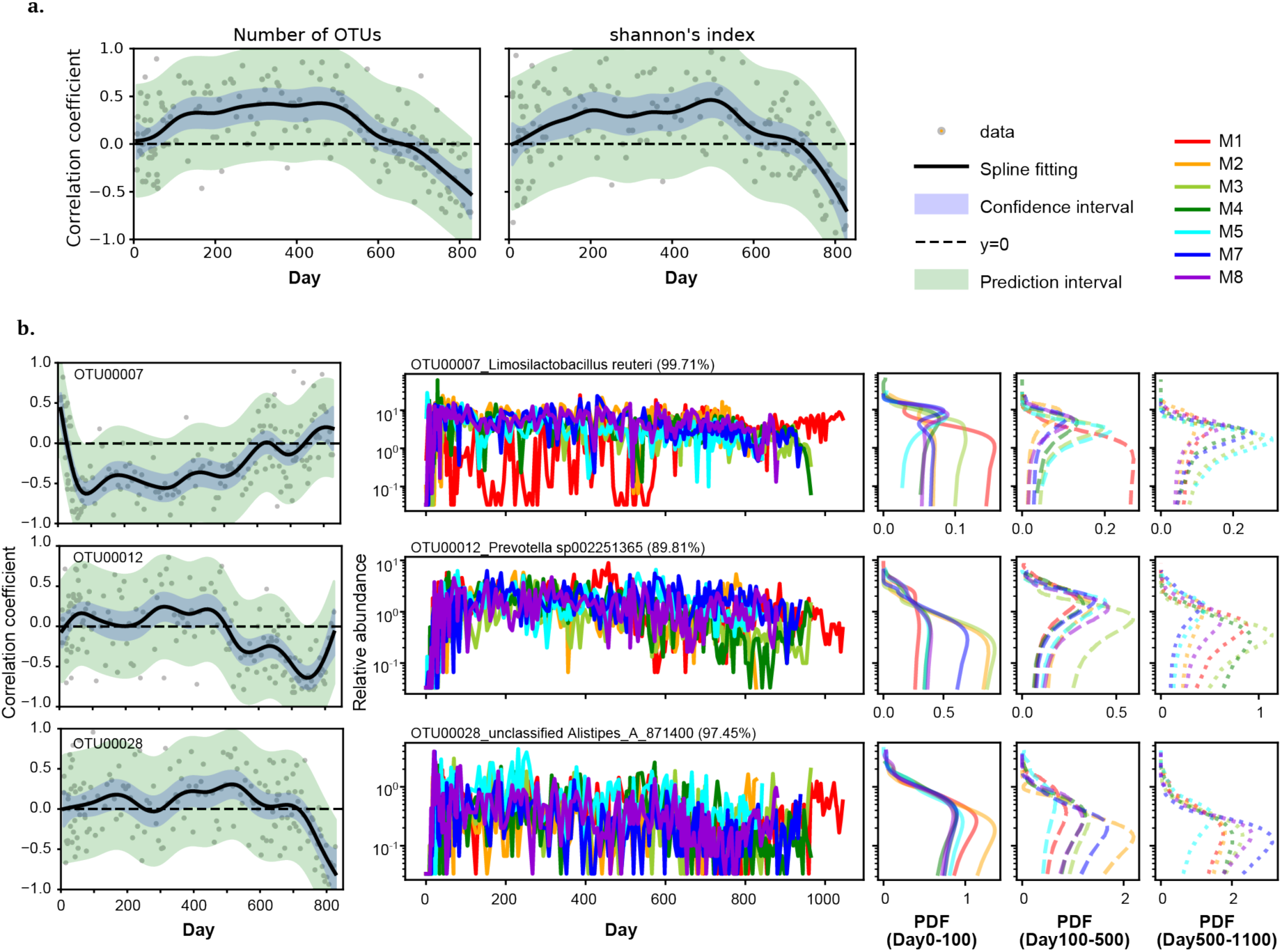
Nonstational inter-individual association with lifespan over time. **a,** Spearman’s correlation coefficients between α-diversities and lifespan. Grey dots indicate Spearman’s correlation coefficient at each time point. Black lines indicate the spline fitting of the coefficients. Blue mesh indicates the confidence interval and green mesh indicates the prediction interval. **b,** Spearman’s correlation coefficients between selected three life-core OTUs and lifespan (left). The annotations for the dots, lines and meshes are the same as **a**. The abundance variation in selected lifecore OTUs (middle) and the abundance probability density function (PDF) for each mouse at each time duration (right).

We also found some of the life-core species showed the non-stationary correlation with lifespan (**Figs. 3b, S11**). OTU00007, which is assigned to *Limosilactobacillus* (*Lactobacillus*) *reuteri,* was negatively correlated with lifespan in early to middle age, though positively correlated in late age. OTU00012, which was most closely aligned to *Prevotella* species (identity 89.81%), showed slightly positive correlation until day 500 though negatively correlated after day 600. OTU00028, closest to *Alistipes* species (identity 97.45%), exhibited a slightly positive correlation until day 700, followed by a negative correlation after day 700.

### OTUs that correlated with life events

Many studies have examined the impact of various events, such as pregnancy, body weight changes, tumorigenesis, and cohousing on the gut microbiome. However, few studies have investigated the relationship between these events and gut microbial changes over the course of an individual’s entire life. In this study, three mice (M3, M7, and Mo) underwent pregnancy and delivery during the study period. Comparison of samples 20 days before pregnancy, during pregnancy, and after delivery revealed significant differences in the unweighted UniFrac distance between all pairs of the three groups. Additionally, significant differences were observed in weighted UniFrac distance between the “before” and “during pregnancy” samples, as well as between the “before” and “after delivery” samples (PERMANOVA, *P* < 0.05; **Fig. S12a**). The number of OTUs did not differ significantly between the samples; however, Shannon’s index was significantly higher in samples collected during pregnancy and after delivery compared to those collected before pregnancy (**Fig. S12b**). The analysis also revealed 59 OTUs with significant changes in abundance between the different sample groups (**Supplementary Table 3**).

We found that 27 OTUs were significantly correlated with body weight, which was only measured in the later life stages, from day 555 to just before death (*P* < 0.05, average rho > 0.3; **Fig. S13**).

Tumors developed in five mice (M2, M6, M7, M8, and Mo) and were detected during autopsies. UniFrac distance analysis of samples collected 30 days before death revealed significant differences in the gut microbiome between mice with and without tumors (**Fig. S14a**). Both the number of OTUs and Shannon’s index were higher in the tumor samples than in the non-tumor samples (**Fig. S14b**). Additionally, we also identified 32 OTUs that were significantly enriched, and 33 OTUs that were significantly depleted, in tumor samples compared to non-tumor samples (**Supplementary Table 4**).

During the experiments, ten mice (eight siblings and parents) were cohoused with different combinations of individuals in different frequencies and periods after initially cohousing all mice together for 51 days after birth (**Fig. 1a**). We divided the samples into two groups: cohoused and non-cohoused, and compared their UniFrac distances. The results revealed that both weighted and unweighted UniFrac distances were significantly lower in the cohoused samples compared to the non-housed samples (**Fig. S15a**). Additionally, the inter-individual distances within a cage were significantly lower after 30 days of cohousing (**Fig. S15b**), suggesting that the microbiomes of individual mice in the same cage became more similar over time.

## Discussion

This is a comprehensive time-course study that elucidates the development of the gut microbiome over the lifespan of cognate SPF mice. The relationship between the gut microbiome and host aging has been well recognized; however, it is based primarily on the comparative analysis of gut microbiomes between individuals of various ages with different dietary habits, lifestyles, and genetic backgrounds(5,23–27). In this study, we used sibling mice and longitudinally monitored changes in the gut microbiome of individuals from birth to death for approximately three years. Although the mice were bred in a controlled environment, phenotypic diversity was ultimately observed among individuals, accompanied by a high inter-individual diversity in the gut microbiomes of the seven siblings that died of natural causes. As C57BL/6J mice were designed as an inbred line and share a common genetic backgrounds, these diverse individual differences are likely caused by stochastic environmental factors.

In the present study, the bimodal distribution indicated that some OTUs can be divided into “life-core” or “transient” OTUs. The “life-core” OTUs were predominantly composed of Bacteroidota, whereas “transient” OTUs were primarily composed of Firmicutes, despite both Bacteroidota and Firmicutes being dominant phyla in gut microbiomes. Interestingly, this result is partially supported on humans by the study conducted by Faith et al(28), which demonstrated long-term stability of Bacteroidota and Actinobacteriota in human gut microbiome. A previous report based on fluorescent in situ hybridization imaging technology showed that Bacteroidota are observed primarily near the intestinal wall, whereas Firmicutes are separated from the gut wall in the mouse intestinal tract(29). These differences in micro-spatial structure within the gut may partially account for the higher persistence of Bacteroidota. In addition to their spatial structure in the gut, Firmicutes have been proposed to be more effective at extracting energy from food than Bacteroidota, thereby promoting more efficient calorie absorption and subsequent weight gain(30,31). Moreover, it is reported that Bacteroidota mainly produce propionate from dietary carbohydrates, whereas Firmicutes produce large amounts of butyrate(32,33). These differences in metabolic capacity might also contribute to the differences in persistence between the two phyla; however, further research is needed to elucidate the precise underlying mechanisms.

Analysis of the variation in OTU abundance revealed 9 distinct patterns across the lifespan of the seven siblings (Fig. 2a). Cluster_4 appeared during early ages, around days 20–50. In particular, dynamic changes in the microbiota were observed during the first 20 days, with Cluster_1 and Cluster_5 emerged on days 14 and 8, respectively (**Fig. S5**). Sequentially, Cluster_7 and Cluster_9 became abundant until day 500–600; Cluster_3 were abundant thereafter. Weighted Jaccard distance analysis revealed that the temporal variations in microbiomes within individuals changed after day ∼500 (**Fig. 1d**). Change–points analysis also suggested certain mode changes related to the gut microbiota dynamics during ages that correspond to early days (day <100) and middle ages (day 300–800). However, we need to consider these results are based on data from a single experiment (**Fig. S6**).

We found a positive correlation in middle age (day 200–500) and a negative correlation in old age (after day 700) between lifespan and α-diversity (Fig. 3a). Given that several diseases have been shown to lower the diversity of gut microbiome(34–36), it is often posited that higher diversity is related to a healthier life. However, in this study, the average α-diversity throughout life was not strongly correlated with lifespan (**Fig. S10**). A tendency for increased gut microbiome diversity was observed shortly before death (**Fig. 1f**), which may have contributed to the reversal in the correlation between α-diversity after day 700 and lifespan. Specifically, the diversity within the last 20 days of life tended to be higher than during the second-to-last 20 days in all mice, except for M1. However, over a longer span, a gradual increase in diversity was observed during the final 100 days of life, even for M1 (**Fig. S2a**). Nevertheless, further studies are needed to verify this phenomenon and determine the time scale over which the increase in diversity occurs before death.

We also found a transition in the correlation patterns of life-core OTUs as well (**Fig. 3b**). *Limosilactobacillus (Lactobacillus)* was negatively correlated with mid-life and exhibited a significant positive correlation in later life (after day 800). The increase in *Lactobacillus* has been reported as associated with lifespan elongation under limited calorie conditions in mice(37). In our study, a consistent tendency was observed in aged mice but not in young mice. The change in correlation tendency with age suggested that the previously reported relationship between microbes and lifespan might be transient under the timescale of lifespan.

We also characterized microbiome changes associated with pregnancy, body weight, tumor development, and cage effect (**Figs. S12–15**). The limited number of mice poses a challenge in accurately determining the influence of these events on the microbiome dynamics and lifespan. However, the observation of gut microbiome changes in before and after the pregnancy, the changes before death associated with tumor development, and time-associated changes according to the cage transfers are intriguing. Although numerous studies have investigated microbiomes in relation to these life events, longitudinal research focusing on dynamic changes within the same individual over time remains limited(38–44). We hope the detailed sampling in this study will provide insight into the characteristic periods associated with among various life events, which may serve as a valuable resource for informing future sampling strategies(45).

In this study, we characterized the life-core/transient microbiome and our result suggest a non-static relationship between the host and the gut microbiome. Further investigation of host immune changes or bacterial functional interactions will help clarify the mechanism(s) underlying the temporal dynamics of diversity and bacterial abundance observed in this study.

## Methods

### Animals and Sample Collections

Two SPF C57BL/6J mice were maintained and bred under the specific pathogen-free conditions at the RIKEN Center for Integrative Medical Sciences Animal Facility (Kanagawa, Japan). Mice were identified using ear punches and housed in rooms maintained at a temperature of 23 ± 2°C and a relative humidity of 50% ± 10%, with a 12-hour light-dark cycle. The animals were housed in cages (Fig. 1a) with wood shavings and provided with food (CE-2, CLEA Japan, Inc.) and water *ad libitum.* The male mouse (M1) was fed Oriental CMF (Oriental Yeast, Tokyo) from days 462 to 559 for breeding. The mouse cages were replaced primarily to control reproduction. The starting point of this study was the birth of the second generation of SPF mice. Nine offspring were born. Male and female offspring were separated after weaning for 50–51 days of age. One of the nine offspring (M9) died at 6 days of age and was therefore excluded from the analysis. Another mouse (M6) was excluded from the analysis of the association between the gut microbiome and host lifespan because it died during retention during sample collection due to debilitation. Mouse feces were collected, snap-frozen, and stored at -80°C. All animal experiments were approved by the Institutional Animal Care and Use Committees of RIKEN Yokohama Branch.

### Autopsies of dead mice

The carcasses of mice that died naturally were immediately stored in a -80°C freezer upon discovery. Autopsies were performed to examine the kidneys, spleen, lungs, heart, cervical lymph nodes, reproductive organs, and liver. If tumors were detected, death was attributed to the tumor; otherwise, it was categorized as age-related. If tumors were detected, death was attributed to the tumor; otherwise, it was categorized as age-related.

### DNA extraction

Bacterial genomic DNA was extracted from mouse feces using the enzymatic lysis method. Frozen fecal pellets were thawed and suspended in 1 mL of TE10 buffer (10 mM Tris-HCl, 10 mM EDTA) containing RNaseA (final concentration of 100 µg/mL; Nippon Gene, Tokyo, Japan). Lysozyme (Sigma-Aldrich; St. Louis, MO, USA) was added at a final concentration of 15 mg/mL, and the suspension was incubated for 1 h at 37 °C with gentle mixing. Purified achromopeptidase (Wako Pure Chemical, Osaka, Japan) was added at a final concentration of 2000 units/mL, and the suspension was further incubated for 30 min at 37 °C. Sodium dodecyl sulfate (final concentration, 1 mg/mL) and proteinase K (final concentration, 1 mg/mL; MERCK Group Japan, Tokyo, Japan) were added to the sample, and the mixture was incubated for 1 h at 55 °C. DNA was extracted with phenol/chloroform/isoamyl alcohol (25:24:1), precipitated with isopropanol and 3 M sodium acetate, washed with 75% ethanol, and resuspended in 200 µL of TE buffer. DNA was purified with a 20% PEG solution (PEG6000 in 2.5 M NaCl), pelleted by centrifugation, rinsed with 75% ethanol, and dissolved in TE buffer.

### 16S rRNA gene amplicon sequencing

We analyzed a total of 51.4 million (51,377,124 reads) high-quality reads of the 16S rRNA gene V1–2 region from eight siblings and the parents (Supplementary Table 1). The isolated DNA (40 ng) was used for PCR amplification of the V1–V2 hypervariable regions of the 16S rRNA gene using the universal primers 27Fmod (5′-AATGATACGGCGACCACCGAGATCTACACxxxxxxxxACACTCTTTCCCTACACGACGCTCTTCC GATCTagrgtttgatymtggctcag-3′) and 338R (5′-CAAGCAGAAGACGGCATACGAGATxxxxxxxxGTGACTGGAGTTCAGACGTGTGCTCTTCCGAT CTtgctgcctcccgtaggagt-3′), containing the Illumina Nextera adapter sequence and an unique 8-bp index sequence (indicated above as xxxxxxxx) for each sample. Thermal cycling was performed on a 9700 PCR system (Life Technologies, Carlsbad, CA, USA) using Ex Taq polymerase (Takara Bio, Tokyo, Japan) with the following cycling conditions: initial denaturation at 96 °C for 2 min; 25 cycles of denaturation at 96 °C for 30 s, annealing at 55° C for 45 s, and extension at 72 °C for 1 min; final extension at 72 °C. All amplicons were purified using AMPure XP magnetic purification beads (Beckman Coulter, Brea, CA, USA) and quantified using the Quant-iT PicoGreen dsDNA Assay kit (Life Technologies). An equal amount of each PCR amplicon was pooled and subjected to sequencing on a MiSeq platform (Illumina, San Diego, CA, USA) using the MiSeq Reagent Kit v3 (600 cycles) according to the manufacturer’s instructions.

### 16S sequence data analysis

After demultiplexing the 16S sequence reads based on a sample-specific index, paired-end reads were joined using the fastq-join program. Reads with average quality values <25 and inexact matches to the universal primer sequences were filtered out. From each sample, 3,000 reads that passed the quality filter were randomly selected for downstream analysis. The selected reads of all the samples were first sorted by the frequency of redundant sequences and grouped into OTUs using UCLUST (https://www.drive5.com/) with a sequence identity threshold of 97%. The taxonomic assignment of each OTU was determined by similarity searching against the Greengenes2 (v2022.10)(21) using the GLSEARCH program (v36.3.8d).

### Temporal variations in β-diversity across a lifetime

To analyze temporal changes in microbial communities, we performed Principal Coordinate Analysis (PCoA) on UniFrac distance data using decomposition.PCA function from sklearn library.

We used the adonis function from the R vegan package (ver 2.6-4) with the 1000 times permutation option for PERMANOVA analysis, and the cmdscale function from the R MASS package for PCoA analysis. We calculated the weighted Jaccard distances between sample A and sample B as follows:

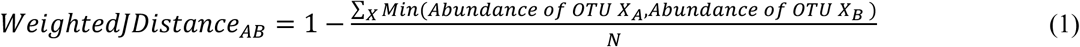

where N indicates the total number of reads in each sample (N = 3,000).

To calculate β-diversity over time, we divided samples into bins every 20 days from day 0, and the average microbial abundance at the OTU level was calculated. To assess the rate of variation of the gut microbiome over the lifetime of an individual, we calculated the weighted Jaccard distance Eq.(1) between the consecutive bins.

### Temporal variations in **α-**diversity across a lifetime

To gain insights into the overall trend and smooth the short-term fluctuations in OTU abundance over time, a simple moving average was calculated for the normalized OTU number. The number of OTU were first normalized with division by a maximum OTU number for each individual throughout the lifespan; the simple moving average for 20 days was then applied using the rolling function in the Pandas library in Python.

### Temporal dynamics clusters analysis

The temporal dynamics clusters were calculated on the time-course data for OTU relative abundances. The Bray–Curtis distances between the OTU abundances were calculated with python scipy.spatial.distance.braycurtis function and hierarchically clustered using the python scipy.cluster.hierarchy.linkage function with method “ward”. Clusters were created using the scipy.cluster.hierarchy.fcluster function with a threshold of 100.0.

### Statistical analyses related to life events and external factors

We conducted the following statistical analyses to investigate changes in the microbiome associated with life events and external factors observed during the experiment, including pregnancy, body weight change, tumorigenesis, and cohousing. All statistical analyses were performed using the R software program (v4.1.1); the wilcox.exact function in the exactRankTests package was used to perform the Wilcoxon rank sum test and the p.adjust function in the R software program (v4.1.1) to perform the Benjamin–Hochberg correction. PERMANOVA compared the gut microbiome structure among groups based on weighted and unweighted UniFrac distances using the adonis function from the R vegan package (ver 2.6-4) with 1000 times permutations. The results were considered statistically significant at *P* < 0.05. OTUs with maximum relative abundance ≥0.1% were used for the following analysis.

For pregnancy, α-diversity indices and the relative abundance of OTUs in the samples collected 20 days before pregnancy (“Before”), 20 days during pregnancy (“During”), and 20 days after delivery (“After”) were compared by the Wilcoxon rank sum test, followed by Benjamin–Hochberg correction. The differences in the gut microbiomes of these three groups were examined by PERMANOVA, with subsequent Benjamin–Hochberg correction.

Meanwhile, for tumorigenesis, α-diversity indices and the relative abundance of OTUs within the samples collected in the last 1 month (30 days) of life for the five mice with tumors detected via autopsy (M2, M6, M7, M8, and Mo) and the non-tumor mice were compared by the Wilcoxon rank sum test. The differences in gut microbiomes between these two groups were examined by PERMANOVA.

To assess the correlation between body weight and OTU abundance, body-weighted information (observation from day 555) and OTU (maximum relative abundance ≥ 0.1%) abundance information was used. Spearman’s correlations were calculated using the spearmanr function in python scipy library.

To evaluate the cohousing cage effect, the weighted/non-weighted UniFrac distances of the mice microbiomes within the same cage and different cages after day 20 were compared by the Wilcoxon rank sum test. The inter-individual weighted UniFrac distances assessed mice in the same cage within the first 30 days after cohousing compared with those after 30 days using the Welch’s *t*-test (ttest_ind function of scipy.stats module in Python 3.10.13).

### Correlation analysis of the gut microbiome with lifespan

To elucidate the trends in the temporal abundance of OTUs and α-diversity at distinct life stages, we computed Spearman’s correlation coefficients at each time point with scipy.stats.spearmanr function. To visualize longitudinal correlation patterns by fitting a smoothing spline through time-indexed correlation data, we applied a generalized additive model (GAM) to the data using the pyGAM package (v0.9.1). The model’s prediction line and confidence intervals were calculated over a defined day range (day 0-850), and confidence intervals were retrieved using gam.confidence_intervals function. Predicted values were calculated with gam.predict function, and residuals were analyzed to construct a 95% prediction interval based on the standard deviation.

### Change–point detection in OTUs

We aim to statistically identify the points where the average abundance changes in the time series(22) . The following algorithm recursively detects multiple change points:

1. For the median of the time series and time, count the number of four events:

- a : the number of *x*(*t*) > *m* for *t* ≤ *t*_*c*_
- b : the number of *x*(*t*) > *m* for *t* > *t*_*c*_
- c : the number of *x*(*t*) ≤ *m* for *t* ≤ *t*_*c*_
- d : the number of *x*(*t*) ≤ *m* for *t* > *t*_*c*_
2. Perform Fisher’s exact test based on the contingency table 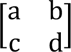. To find a significant change point, search for 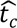 which has the smallest p-value.

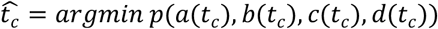
3. If the minimum p-value is less than a certain significance level, then the boundary 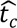, is regarded as a change–point.
4. Perform the above operations recursively on the time series before and after the division by 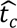.
5. If the minimum p-value exceeds α, terminate the process.

## Data availability

The 16S rRNA V1–V2 region sequences analyzed in the current study were deposited in the DDBJ/GenBank/EMBL repositories with the accession code DRA015228.

## Acknowledgments

We thank K. Kaida, C. Shindo, J. Noack, M. Takagi, and M. Tanokura (RIKEN) for their technical support. We are also supported by A. Siefert from the Cornell CSCU department for statistical advices. This work was funded by JSPS KAKENHI (Grant Number 21409803) to L.T., and a RIKEN Integrated Symbiology grant to M.H. and W.S., and JST, CREST Grant Number JPMJCR22N3, Japan to L.T, M.T. and W.S..

## Author contributions

L.T. contributed toward the experimental design, sampling and experiments, statistical analysis, writing of the original draft, reviewing and editing of the manuscript, and funding acquisition; E.W. and T.U. contributed toward sampling and experiments; R.M. contributed toward changing-point analysis; S.B. contributed toward statistical analysis; R.K., Y.O., Y.K., and H.M., contributed toward sequencing data processing and ; M.U., H.T., M.T. contributed toward reviewing and editing of the manuscript; M.H. contributed toward reviewing and editing of the manuscript and funding acquisition; W.S. contributed toward the experimental design, reviewing and editing of the manuscript, supervision, and funding acquisition.

## References

1. Gordon HA, Bruckner-Kardoss E, Wostmann BS. Aging in germ-free mice: life tables and lesions observed at natural death. J Gerontol. 1966;21(3):380–7.

2. Zhang C, Li S, Yang L, Huang P, Li W, Wang S, et al. Structural modulation of gut microbiota in life-long calorie-restricted mice. Nat Commun. 2013;4(1):2163.

3. Conlon MA, Bird AR. The Impact of Diet and Lifestyle on Gut Microbiota and Human Health. Nutrients. 2015 Jan;7(1):17–44.

4. Dogra SK, Chung CK, Wang D, Sakwinska O, Colombo Mottaz S, Sprenger N. Nurturing the Early Life Gut Microbiome and Immune Maturation for Long Term Health. Microorganisms. 2021 Oct;9(10):2110.

5. Yatsunenko T, Rey FE, Manary MJ, Trehan I, Dominguez-Bello MG, Contreras M, et al. Human gut microbiome viewed across age and geography. nature. 2012;486(7402):222–7.

6. Odamaki T, Kato K, Sugahara H, Hashikura N, Takahashi S, Xiao J zhong, et al. Age-related changes in gut microbiota composition from newborn to centenarian: a cross-sectional study. BMC Microbiol. 2016 Dec;16(1):90.

7. Claesson MJ, Cusack S, O’Sullivan O, Greene-Diniz R, De Weerd H, Flannery E, et al. Composition, variability, and temporal stability of the intestinal microbiota of the elderly. Proc Natl Acad Sci. 2011 Mar 15;108(supplement_1):4586–91.

8. Claesson MJ, Jeffery IB, Conde S, Power SE, O’connor EM, Cusack S, et al. Gut microbiota composition correlates with diet and health in the elderly. Nature. 2012;488(7410):178–84.

9. Biagi E, Franceschi C, Rampelli S, Severgnini M, Ostan R, Turroni S, et al. Gut microbiota and extreme longevity. Curr Biol. 2016;26(11):1480–5.

10. Wu L, Zeng T, Zinellu A, Rubino S, Kelvin DJ, Carru C. A Cross-Sectional Study of Compositional and Functional Profiles of Gut Microbiota in Sardinian Centenarians. Thaiss CA, editor. mSystems. 2019 Aug 27;4(4):e00325–19.

11. Rampelli S, Soverini M, D’Amico F, Barone M, Tavella T, Monti D, et al. Shotgun Metagenomics of Gut Microbiota in Humans with up to Extreme Longevity and the Increasing Role of Xenobiotic Degradation. David LA, editor. mSystems. 2020 Apr 28;5(2):e00124–20.

12. Si J, Vázquez-Castellanos JF, Gregory AC, Decommer L, Rymenans L, Proost S, et al. Long-term life history predicts current gut microbiome in a population-based cohort study. Nat Aging. 2022 Oct;2(10):885–95.

13. Vandeputte D, De Commer L, Tito RY, Kathagen G, Sabino J, Vermeire S, et al. Temporal variability in quantitative human gut microbiome profiles and implications for clinical research. Nat Commun. 2021 Nov 18;12(1):6740.

14. Flores GE, Caporaso JG, Henley JB, Rideout JR, Domogala D, Chase J, et al. Temporal variability is a personalized feature of the human microbiome. Genome Biol. 2014 Dec 3;15(12):531.

15. Voigt AY, Costea PI, Kultima JR, Li SS, Zeller G, Sunagawa S, et al. Temporal and technical variability of human gut metagenomes. Genome Biol. 2015 Apr 8;16(1):73.

16. Stewart CJ, Ajami NJ, O’Brien JL, Hutchinson DS, Smith DP, Wong MC, et al. Temporal development of the gut microbiome in early childhood from the TEDDY study. Nature. 2018 Oct;562(7728):583–8.

17. Jeffery IB, Lynch DB, O’Toole PW. Composition and temporal stability of the gut microbiota in older persons. ISME J. 2016 Jan;10(1):170–82.

18. Derrien M, Alvarez AS, Vos WM de. The Gut Microbiota in the First Decade of Life. Trends Microbiol. 2019 Dec 1;27(12):997–1010.

19. RodrÍguez JM, Murphy K, Stanton C, Ross RP, Kober OI, Juge N, et al. The composition of the gut microbiota throughout life, with an emphasis on early life. Microb Ecol Health Dis. 2015 Dec 1;26(1):26050.

20. Greenhalgh K, Meyer KM, Aagaard KM, Wilmes P. The human gut microbiome in health: establishment and resilience of microbiota over a lifetime. Environ Microbiol. 2016;18(7):2103– 16.

21. McDonald D, Jiang Y, Balaban M, Cantrell K, Zhu Q, Gonzalez A, et al. Greengenes2 unifies microbial data in a single reference tree. Nat Biotechnol. 2024 May;42(5):715–8.

22. Sato AH, Takayasu H. Segmentation procedure based on Fisher’s exact test and its application to foreign exchange rates [Internet]. arXiv; 2013 [cited 2024 Dec 31]. Available from: http://arxiv.org/abs/1309.0602

23. Bosco N, Noti M. The aging gut microbiome and its impact on host immunity. Genes Immun. 2021 Oct;22(5):289–303.

24. Ross FC, Patangia D, Grimaud G, Lavelle A, Dempsey EM, Ross RP, et al. The interplay between diet and the gut microbiome: implications for health and disease. Nat Rev Microbiol. 2024 Nov;22(11):671–86.

25. Tamayo M, Olivares M, Ruas-Madiedo P, Margolles A, Espín JC, Medina I, et al. How Diet and Lifestyle Can Fine-Tune Gut Microbiomes for Healthy Aging. Annu Rev Food Sci Technol. 2024 Jun 28;15(Volume 15, 2024):283–305.

26. Hall AB, Tolonen AC, Xavier RJ. Human genetic variation and the gut microbiome in disease. Nat Rev Genet. 2017 Nov;18(11):690–9.

27. Wolter M, Grant ET, Boudaud M, Steimle A, Pereira GV, Martens EC, et al. Leveraging diet to engineer the gut microbiome. Nat Rev Gastroenterol Hepatol. 2021 Dec;18(12):885–902.

28. Faith JJ, Guruge JL, Charbonneau M, Subramanian S, Seedorf H, Goodman AL, et al. The Long-Term Stability of the Human Gut Microbiota. Science. 2013 Jul 5;341(6141):1237439.

29. Earle KA, Billings G, Sigal M, Lichtman JS, Hansson GC, Elias JE, et al. Quantitative Imaging of Gut Microbiota Spatial Organization. Cell Host Microbe. 2015 Oct 14;18(4):478–88.

30. Krajmalnik-Brown R, Ilhan Z, Kang D, DiBaise JK. Effects of Gut Microbes on Nutrient Absorption and Energy Regulation. Nutr Clin Pract. 2012 Apr;27(2):201–14.

31. Magne F, Gotteland M, Gauthier L, Zazueta A, Pesoa S, Navarrete P, et al. The firmicutes/bacteroidetes ratio: a relevant marker of gut dysbiosis in obese patients? Nutrients. 2020;12(5):1474.

32. Louis P, Flint HJ. Formation of propionate and butyrate by the human colonic microbiota. Environ Microbiol. 2017 Jan;19(1):29–41.

33. Manson JM, Rauch M, Gilmore MS. The Commensal Microbiology of the Gastrointestinal Tract. In: Huffnagle GB, Noverr MC, editors. GI Microbiota and Regulation of the Immune System [Internet]. New York, NY: Springer New York; 2008 [cited 2024 Dec 29]. p. 15–28. (Advances in Experimental Medicine and Biology; vol. 635). Available from: http://link.springer.com/10.1007/978-0-387-09550-9_2

34. Kriss M, Hazleton KZ, Nusbacher NM, Martin CG, Lozupone CA. Low diversity gut microbiota dysbiosis: drivers, functional implications and recovery. Curr Opin Microbiol. 2018 Aug;44:34– 40.

35. Brown K, DeCoffe D, Molcan E, Gibson DL. Diet-induced dysbiosis of the intestinal microbiota and the effects on immunity and disease. Nutrients. 2012 Aug;4(8):1095–119.

36. Mosca A, Leclerc M, Hugot JP. Gut Microbiota Diversity and Human Diseases: Should We Reintroduce Key Predators in Our Ecosystem? Front Microbiol. 2016;7:455.

37. Zhang C, Li S, Yang L, Huang P, Li W, Wang S, et al. Structural modulation of gut microbiota in life-long calorie-restricted mice. Nat Commun. 2013 Jul 16;4(1):2163.

38. Lyu X, Wang S, Zhong J, Cai L, Zheng Y, Zhou Y, et al. Gut microbiome interacts with pregnancy hormone metabolites in gestational diabetes mellitus. Front Microbiol. 2023 Jul 10;14:1175065.

39. DiGiulio DB, Callahan BJ, McMurdie PJ, Costello EK, Lyell DJ, Robaczewska A, et al. Temporal and spatial variation of the human microbiota during pregnancy. Proc Natl Acad Sci U S A. 2015 Sep;112(35):11060–5.

40. Roje B, Zhang B, Mastrorilli E, Kovačić A, Sušak L, Ljubenkov I, et al. Gut microbiota carcinogen metabolism causes distal tissue tumours. Nature. 2024 Aug;632(8027):1137–44.

41. Shi Y, Li X, Zhang J. Systematic review on the role of the gut microbiota in tumors and their treatment. Front Endocrinol [Internet]. 2024 Aug 8 [cited 2024 Dec 31];15. Available from: https://www.frontiersin.org/journals/endocrinology/articles/10.3389/fendo.2024.1355387/full

42. Lipinski JH, Zhou X, Gurczynski SJ, Erb-Downward JR, Dickson RP, Huffnagle GB, et al. Cage Environment Regulates Gut Microbiota Independent of Toll-Like Receptors. Infect Immun [Internet]. 2021 Aug 16 [cited 2024 Dec 31];89(9). Available from: https://journals.asm.org/doi/10.1128/iai.00187-21

43. Nemoto S, Kubota T, Ohno H. Exploring body weight-influencing gut microbiota by elucidating the association with diet and host gene expression. Sci Rep. 2023 Apr 5;13(1):5593.

44. Sinha T, Brushett S, Prins J, Zhernakova A. The maternal gut microbiome during pregnancy and its role in maternal and infant health. Curr Opin Microbiol. 2023 Aug 1;74:102309.

45. Karwowska Z, Szczerbiak P, Kosciolek T. Microbiome time series data reveal predictable patterns of change. Microbiol Spectr. 2024 Oct;12(10):e0410923.

