## Supplementary material for "High-resolution temporal profiling of the gut microbiome through the whole life of mice": Fig. S

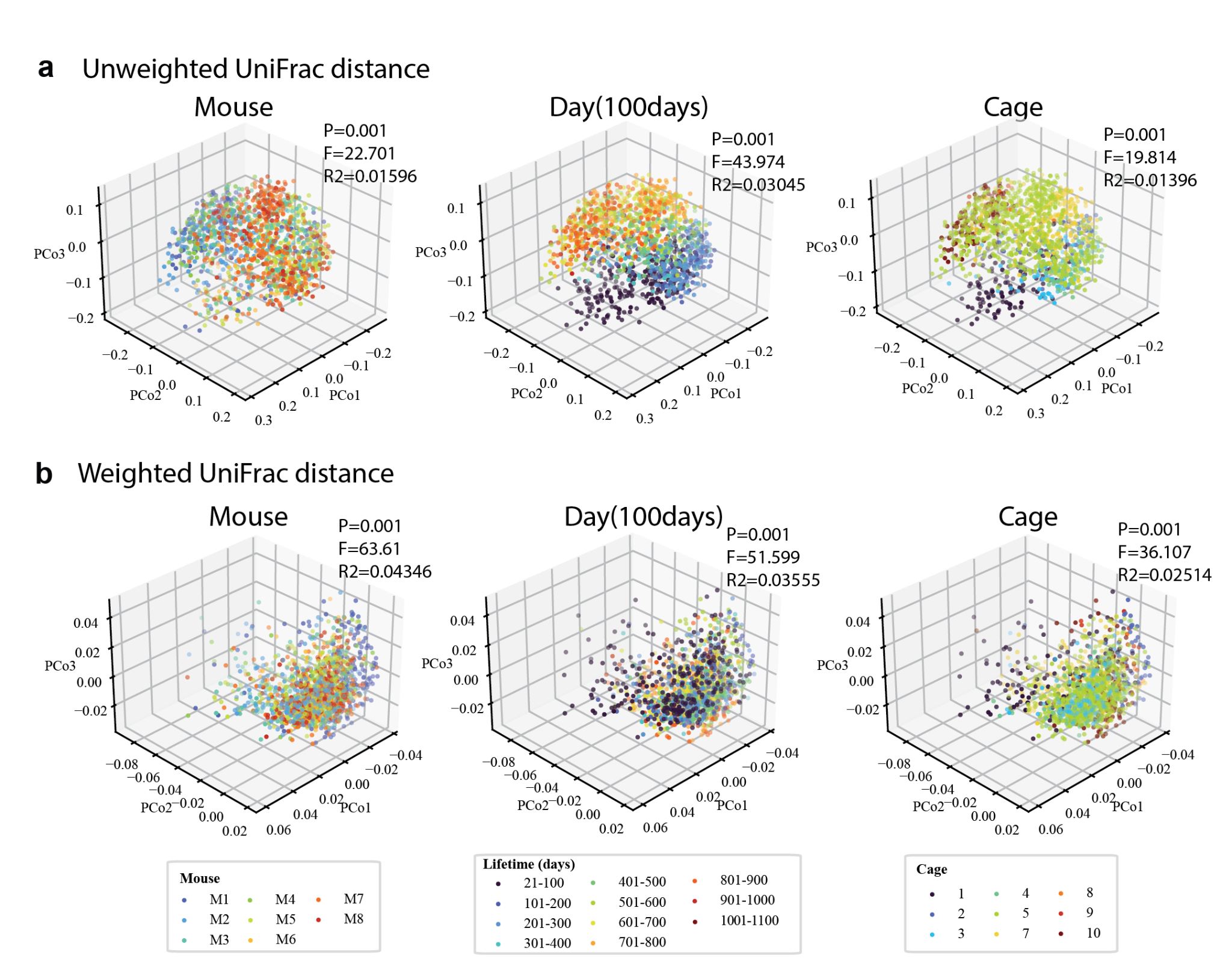

**Supplementary Fig. 1.** Principle coordinate analysis (PCoA) based on UniFrac distance. **a,** PCoA analysis of seven mice (day 20 to day 1044) based on the weighted UniFrac distance. Colored by individual mice (left), the days after birth (center), cage (right), respectively. **b,** PCoA analysis of seven mice (day 20 to day 1044) based on the unweighted UniFrac distance. Colored by individual mice (left), the days after birth (center), cage (right), respectively.

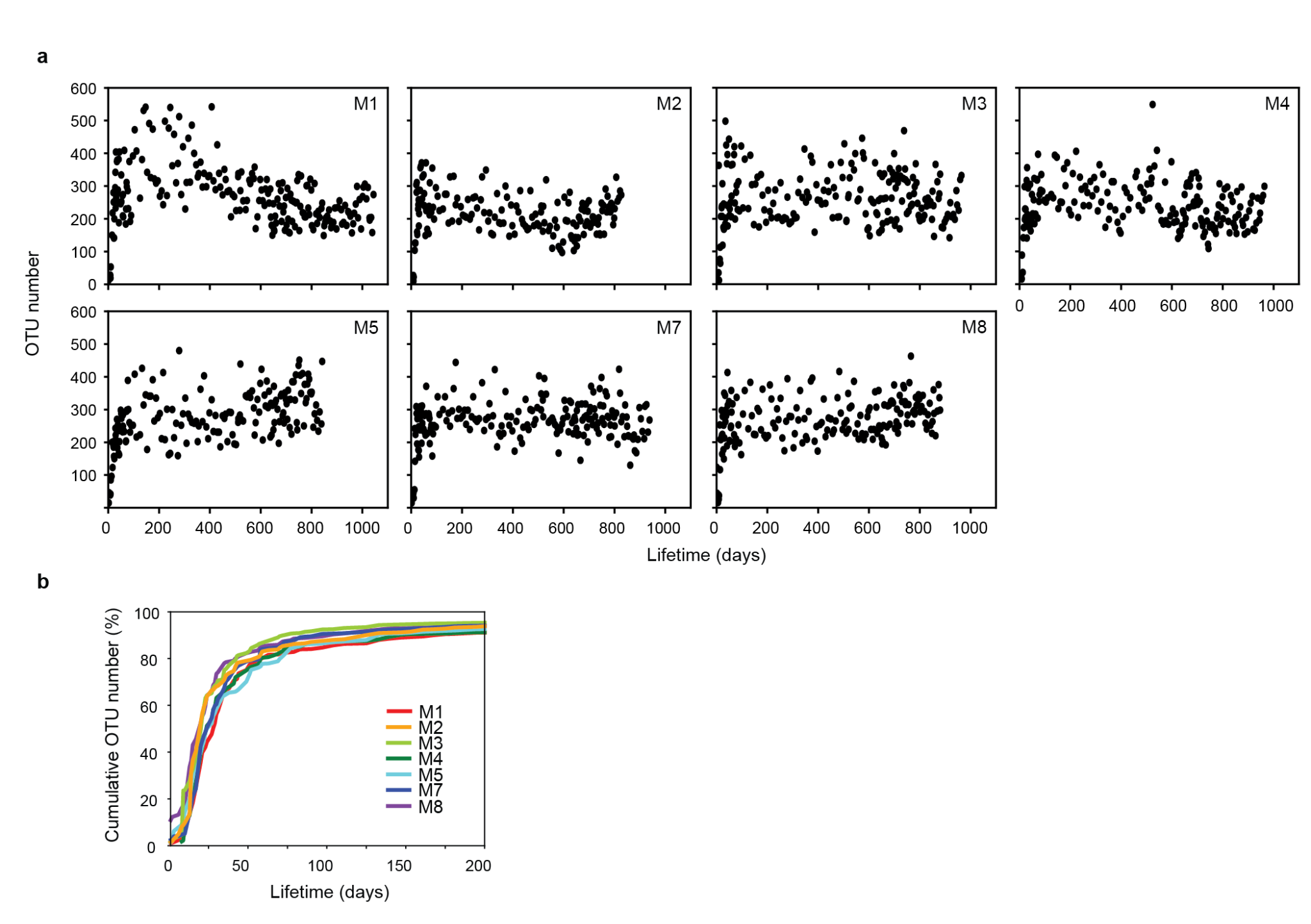
**Supplementary Fig. 2.** Temporal changes of $\alpha$-diversities. **a.** Temporal transition of the number of operational taxonomic units (OTUs). Plot showing the number of OTUs throughout the lifetime of the seven siblings. **b.** Cumulative OTU number over time (%); cumulative OTU number among all OTUs encountered in the lifespan of mice normalized by the total OTU number throughout their life.

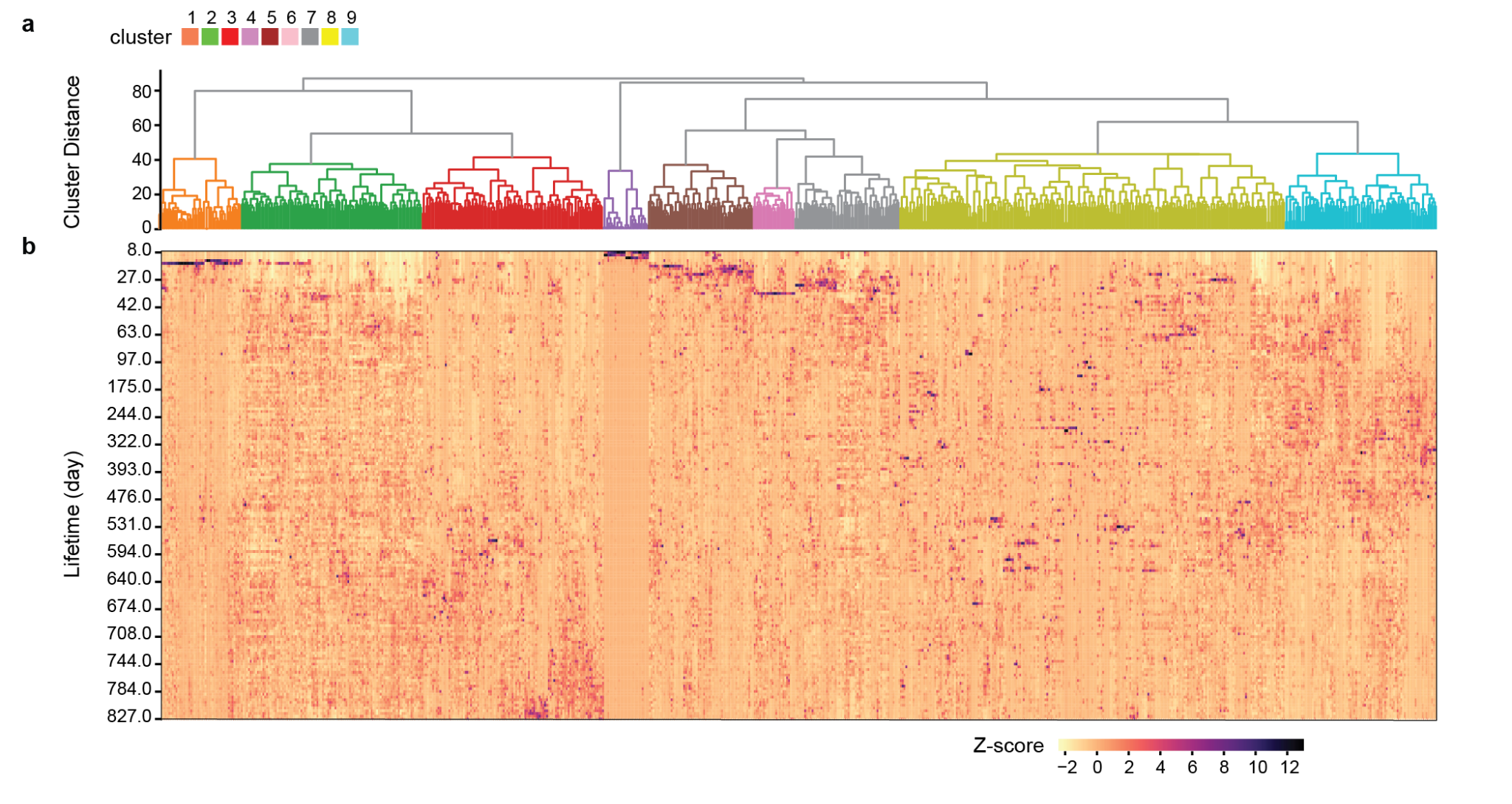

**Supplementary Fig. 3.** Heatmap of the OTUs abundances along time. **a.** Hierarchical clustering analysis (Ward method, Euclidean distance) of the temporal dynamics of OTUs (maximum relative abundance ≥ 0.1%). **b.** Heatmap of the z-scored temporal dynamics of major OTUs.

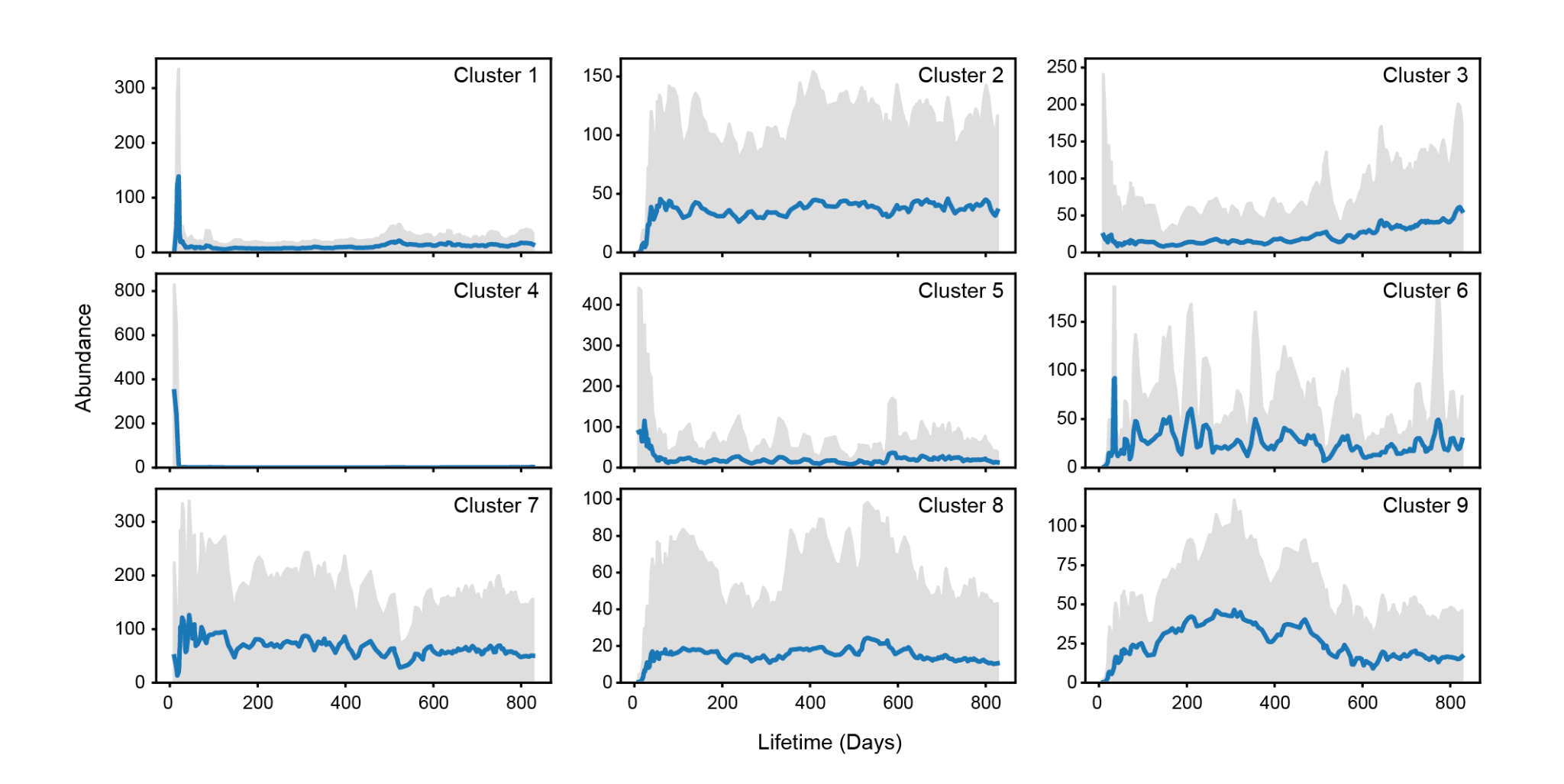

**Supplementary Fig. 4.** Nine Clusters of temporal operational taxonomic unit (OTU) dynamics. The average relative abundance of the OTUs included in each temporal dynamic cluster were determined.

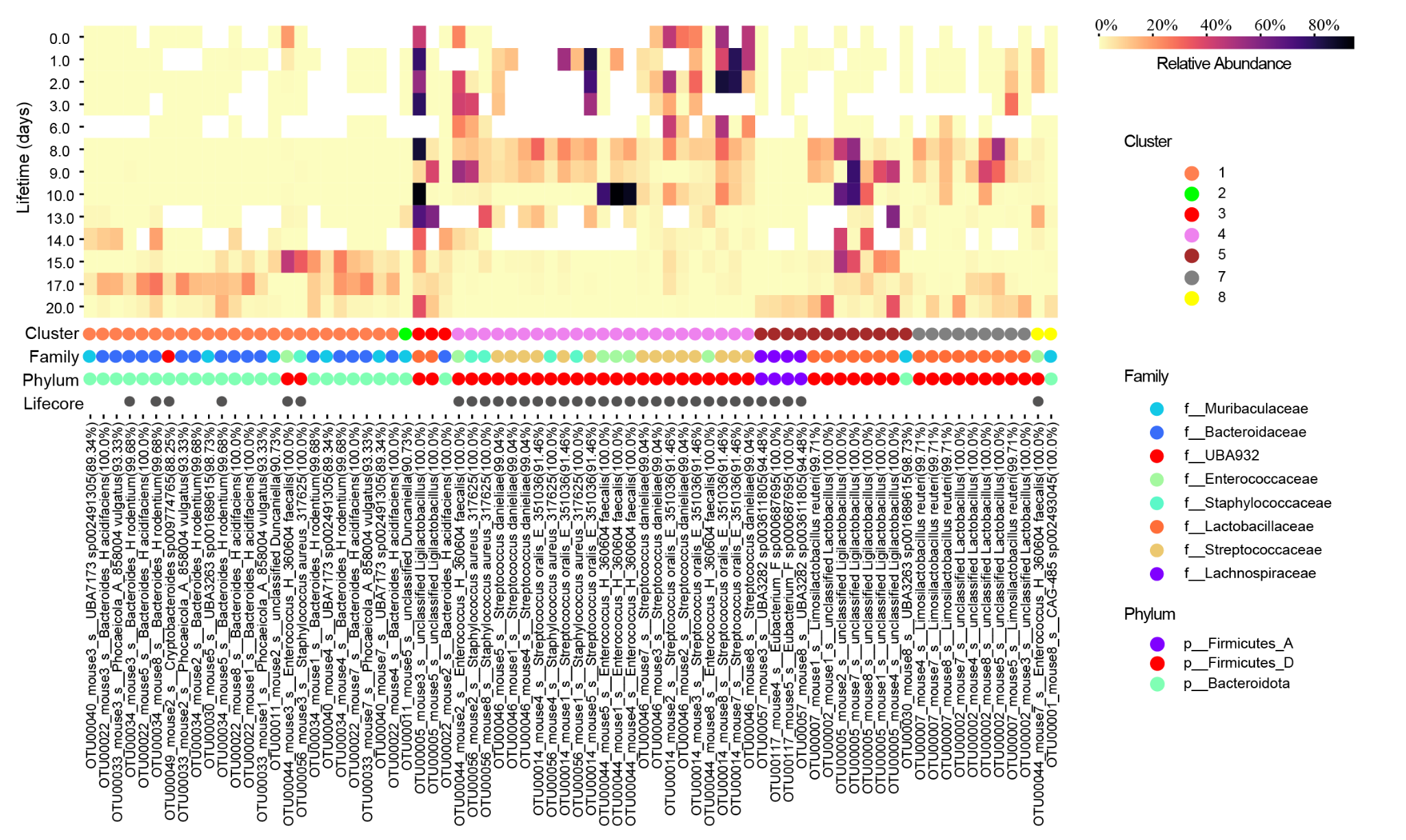

**Supplementary Fig. 5.** Heatmap of the major OTU abundances in early life. Heatmap of the relative abundances of major OTUs (max relative abundance>5%) in the first 20 days of life.

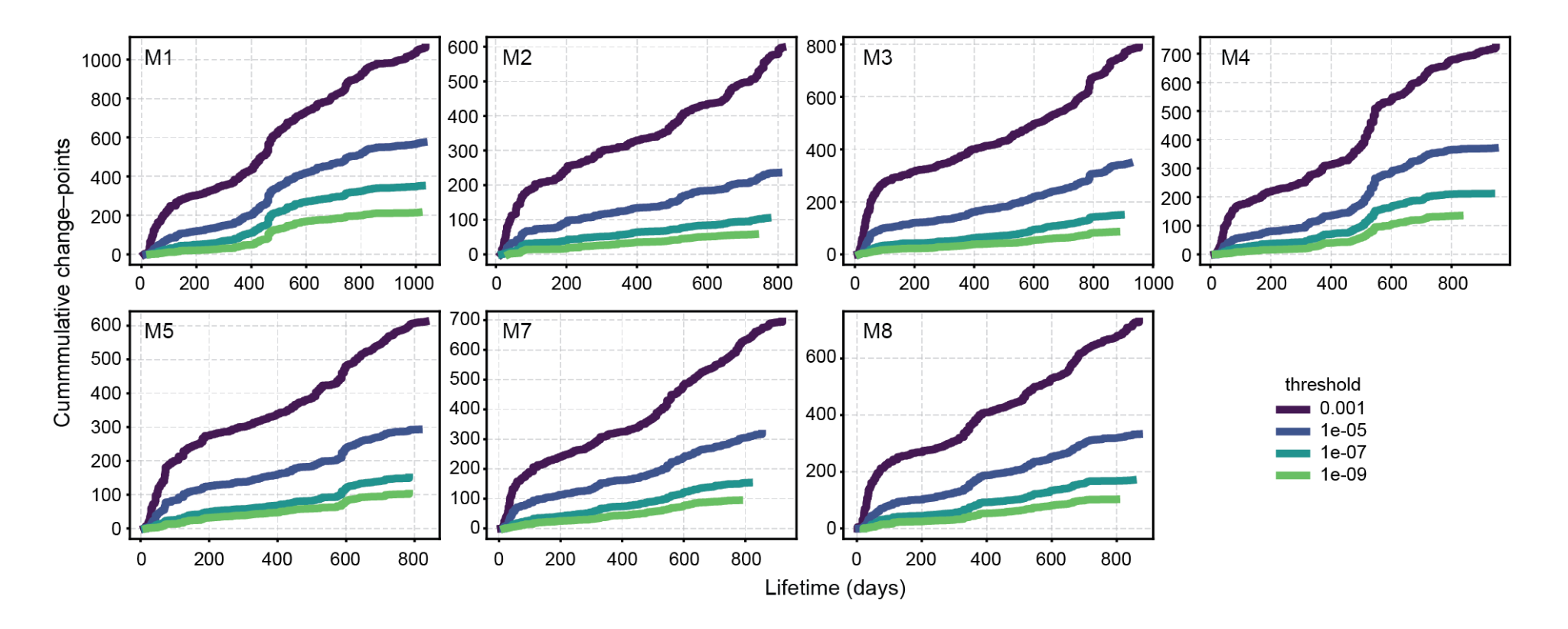

**Supplementary Fig. 6.** Cumulative change–points of OTUs. Each color represents different thresholds in Fisher’s exact test. (See method)

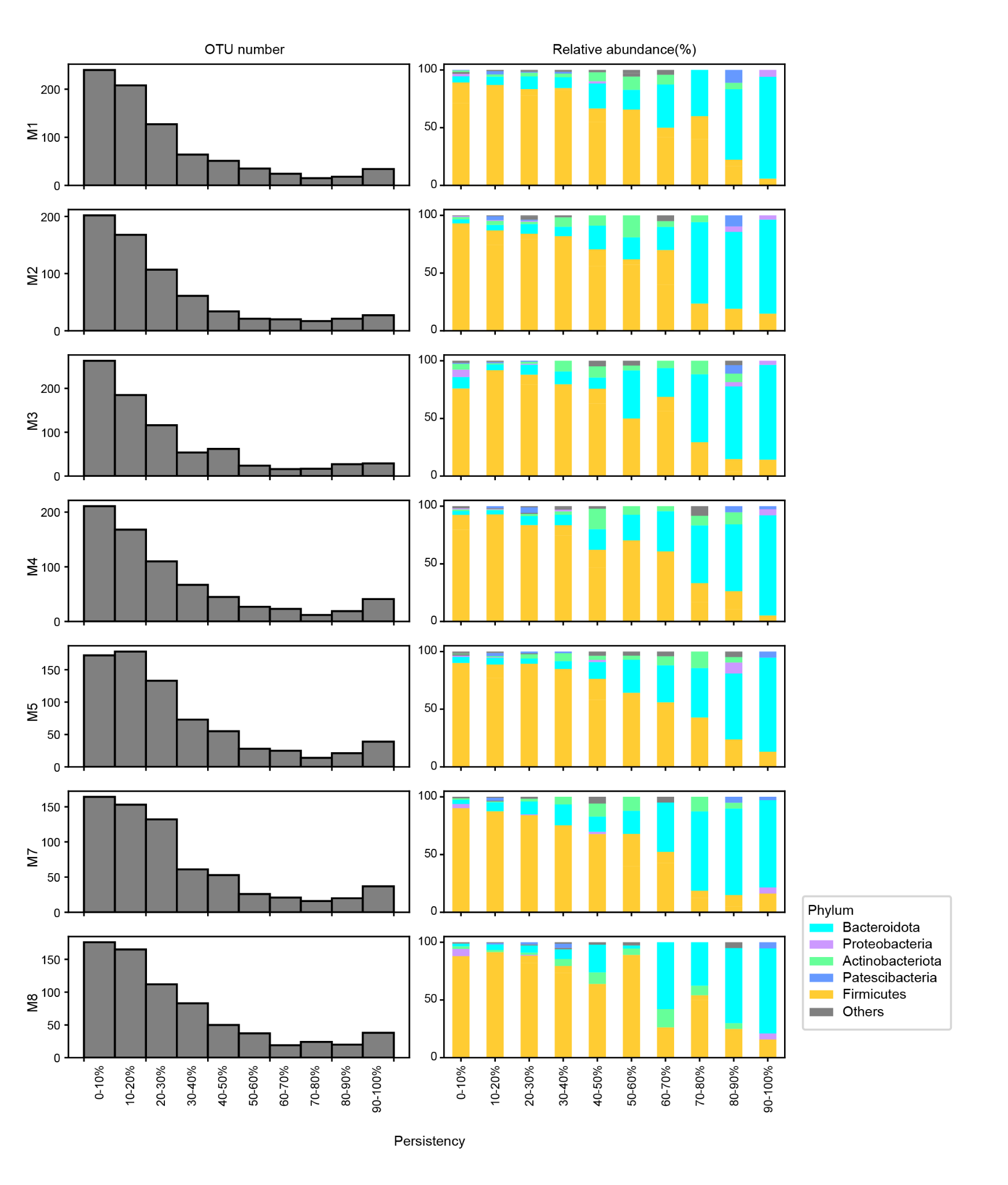
**Supplementary Fig. 7.** Life-core and transient microbiomes. **a.** Histograms of the observed frequency of the operational taxonomic units (OTUs) (maximum relative abundance ≥ 0.1%). The observed frequency of each OTU was calculated by dividing the number of non-zero value counts by the total number of observations. **b.** Bar graphs indicating the phylum-level composition of the OTUs in each bin.

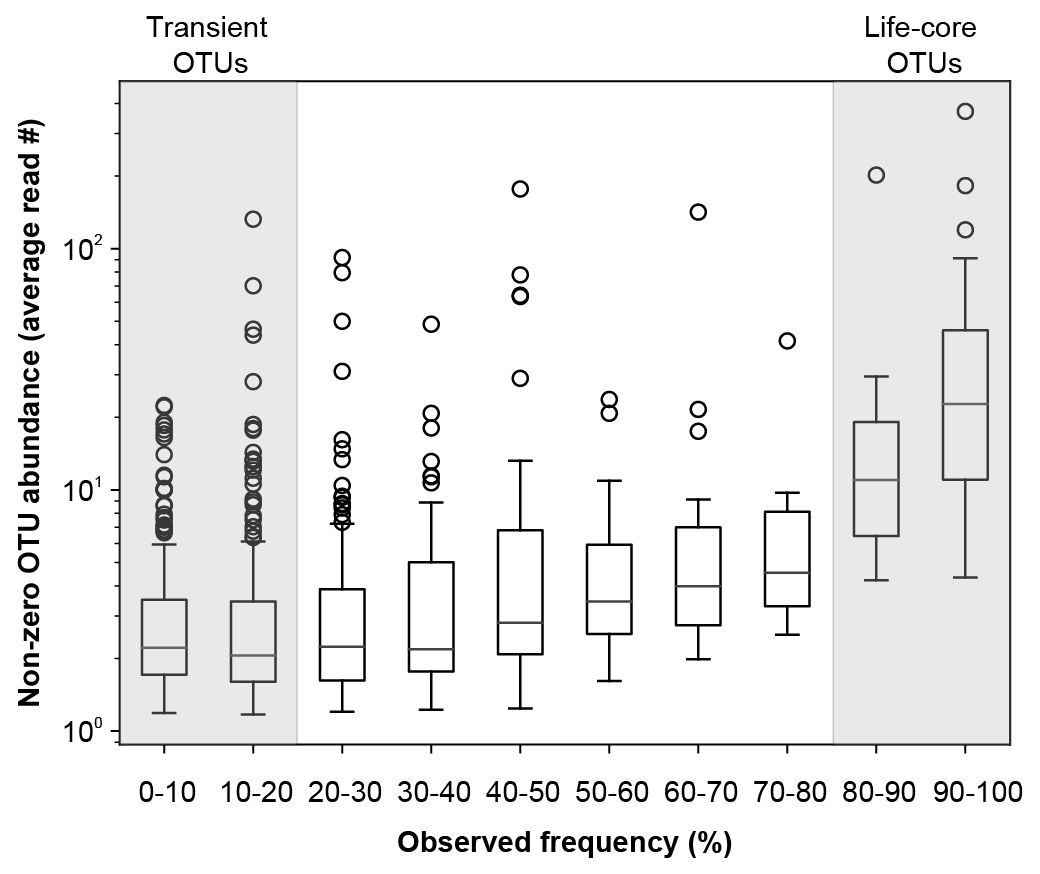
**Supplementary Fig. 8.** Observed frequency and abundance of OTUs. Boxplot of the average abundance of OTUs (maximum relative abundance ≥ 0.1%) when only considering non-zero reads according to the OTUs’ observation frequencies.

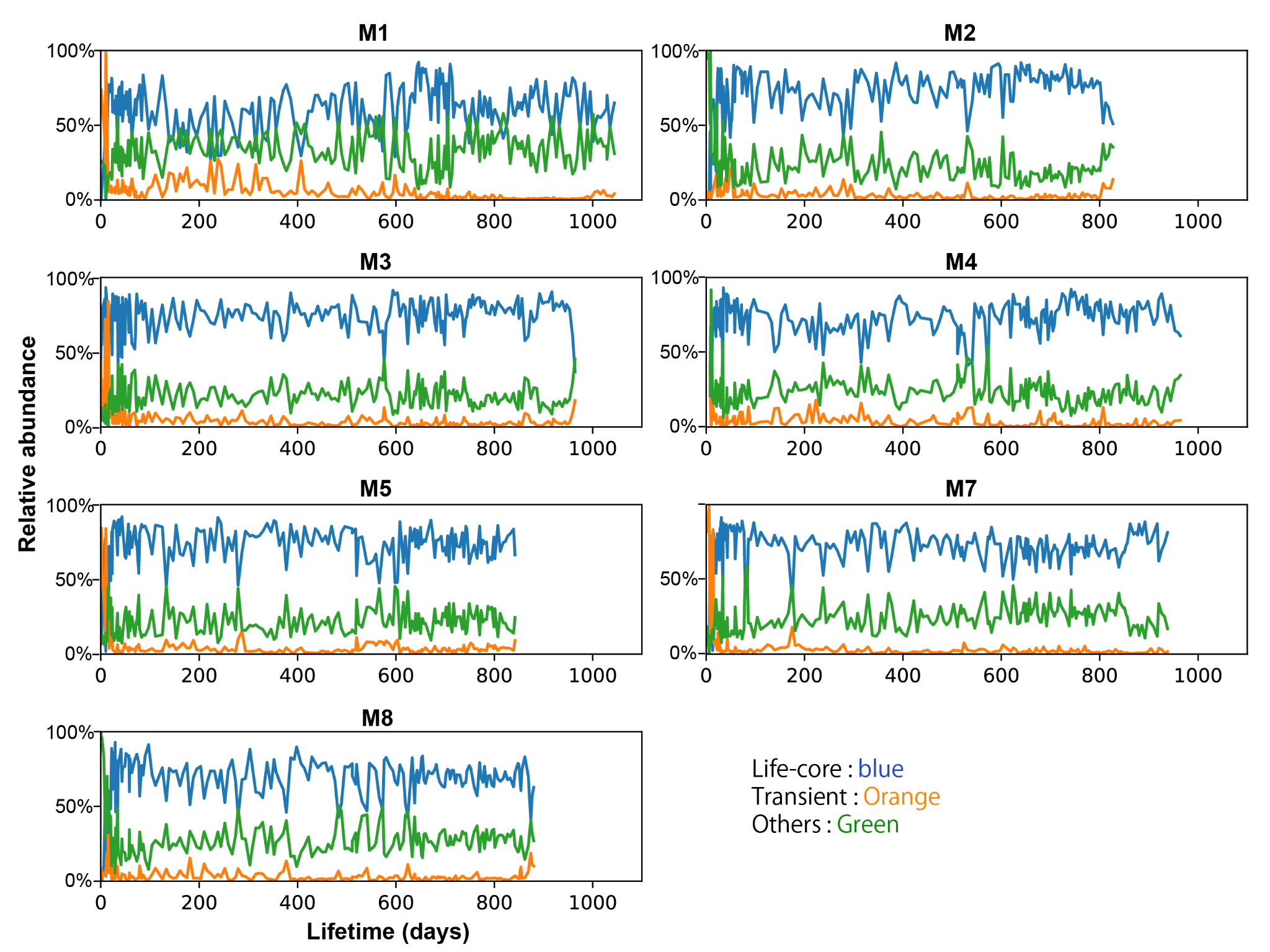

**Supplementary Fig. 9.** Total abundance of life-core/transient OTUs over time. Transition of the total relative abundance of life-core, transient, other OTUs over time.

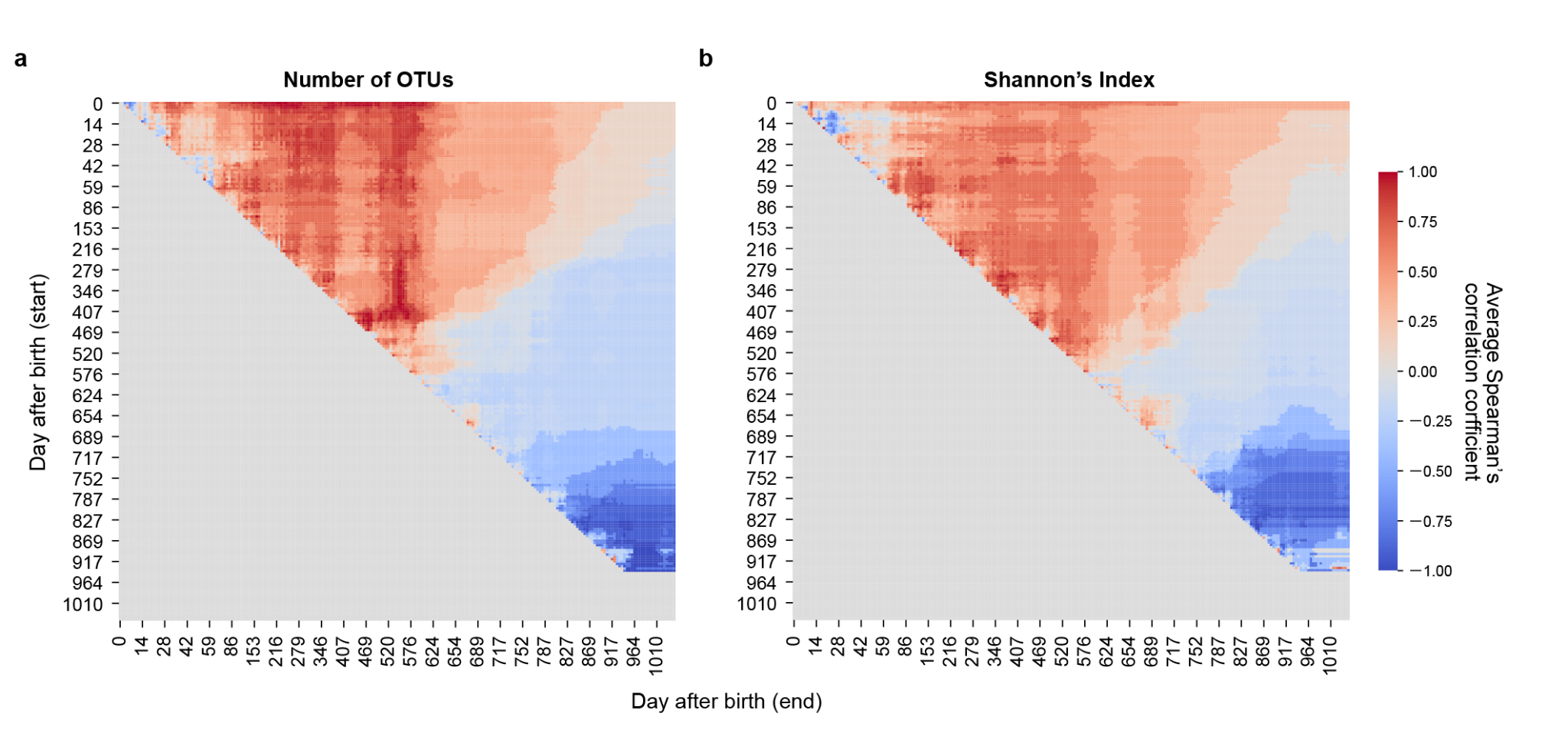

**Supplementary Fig. 10.** Non-stationary correlation between lifespan and $\alpha$-diversity. The Heatmap of Spearman’s rho values between average $\alpha$-diversity and lifespan across all period combinations. The vertical axis denotes the first day of the diversity calculation period, and the horizontal axis represents the last day of the period. e.g., the point at the top right corner signifies the correlation coefficient between the average diversity spanning all time points from day 0 to the last observation point**.**

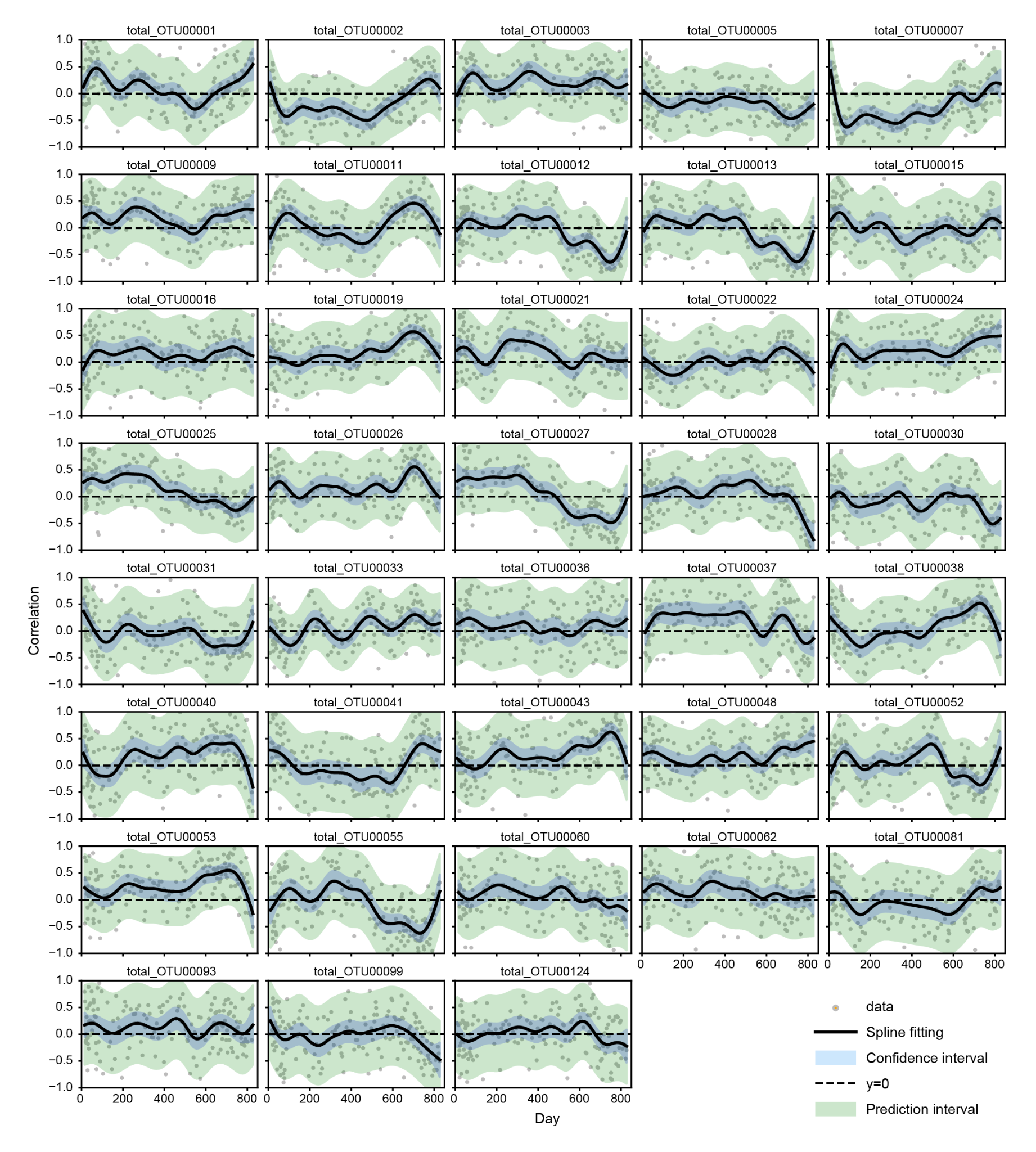

**Supplementary Fig. 11.** Correlation of life-core OTU abundances and lifespan in mice. Spearman’s rho values between the abundance of life-core OTUs and lifespans. Grey dots indicate Spearman’s correlation coefficient at each time point. Black lines indicate the spline fitting of the coefficients. Blue mesh indicates the confidence interval and green mesh indicates the prediction interval.

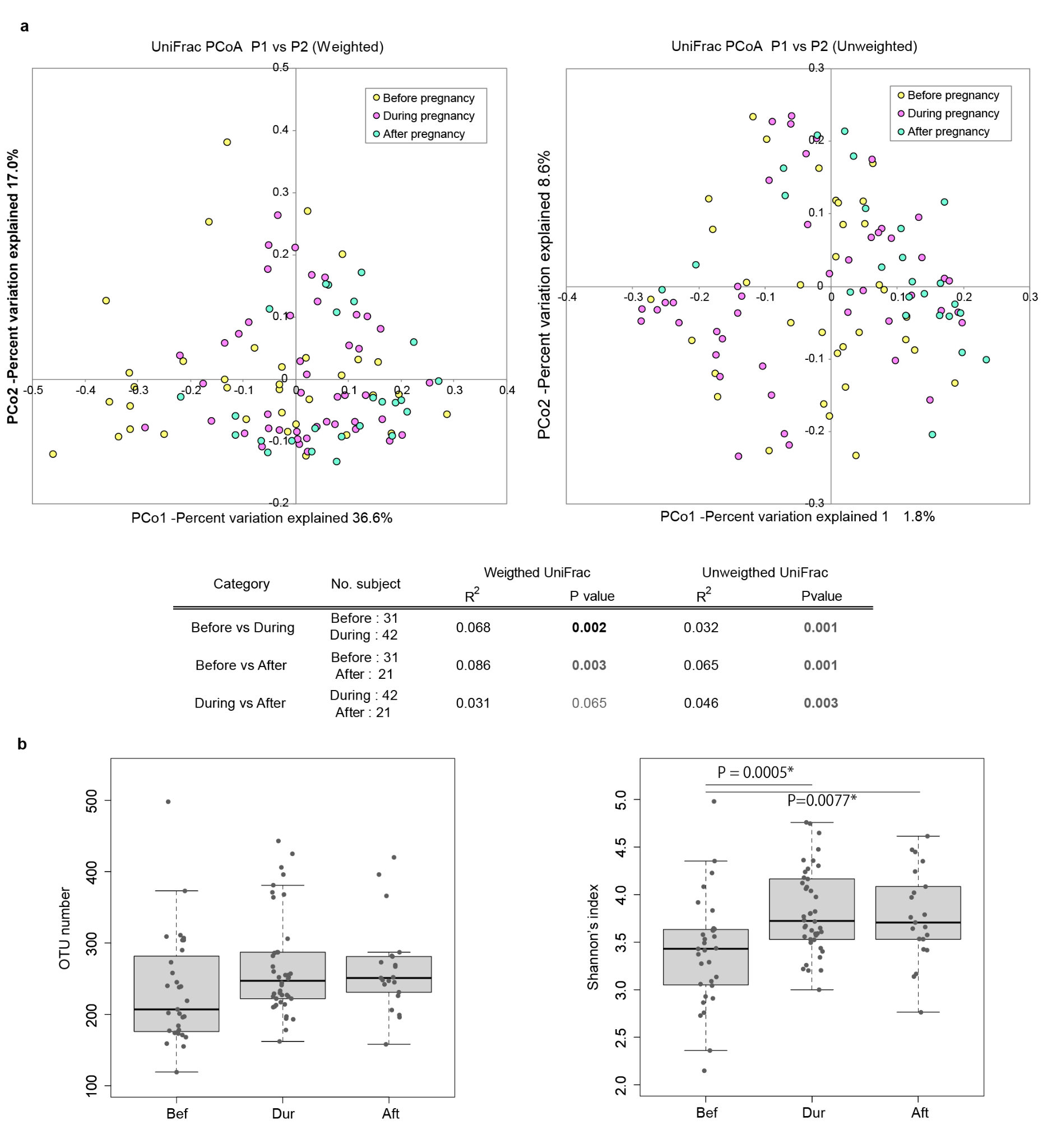

**Supplementary Fig. 12.** The effect of pregnancy. **a.** Principal coordinate analysis (PCoA) of the microbiome composition 20 days before pregnancy to 20 days after delivery based on weighted/unweighted UniFrac distances. The period up to 20 days before the date of delivery was considered “during pregnancy.” Permutational multivariate analysis of variance (PERMANOVA) results between the three groups (before pregnancy, during pregnancy, and after pregnancy) are shown below. The pre-pregnancy period, coinciding with another birth period was excluded from analysis. **b.** Boxplot of the number of OTUs and Shannon’s index in the three groups (Wilcoxon rank sum test with Benjamin–Hochberg correction, * *P* < 0.05).

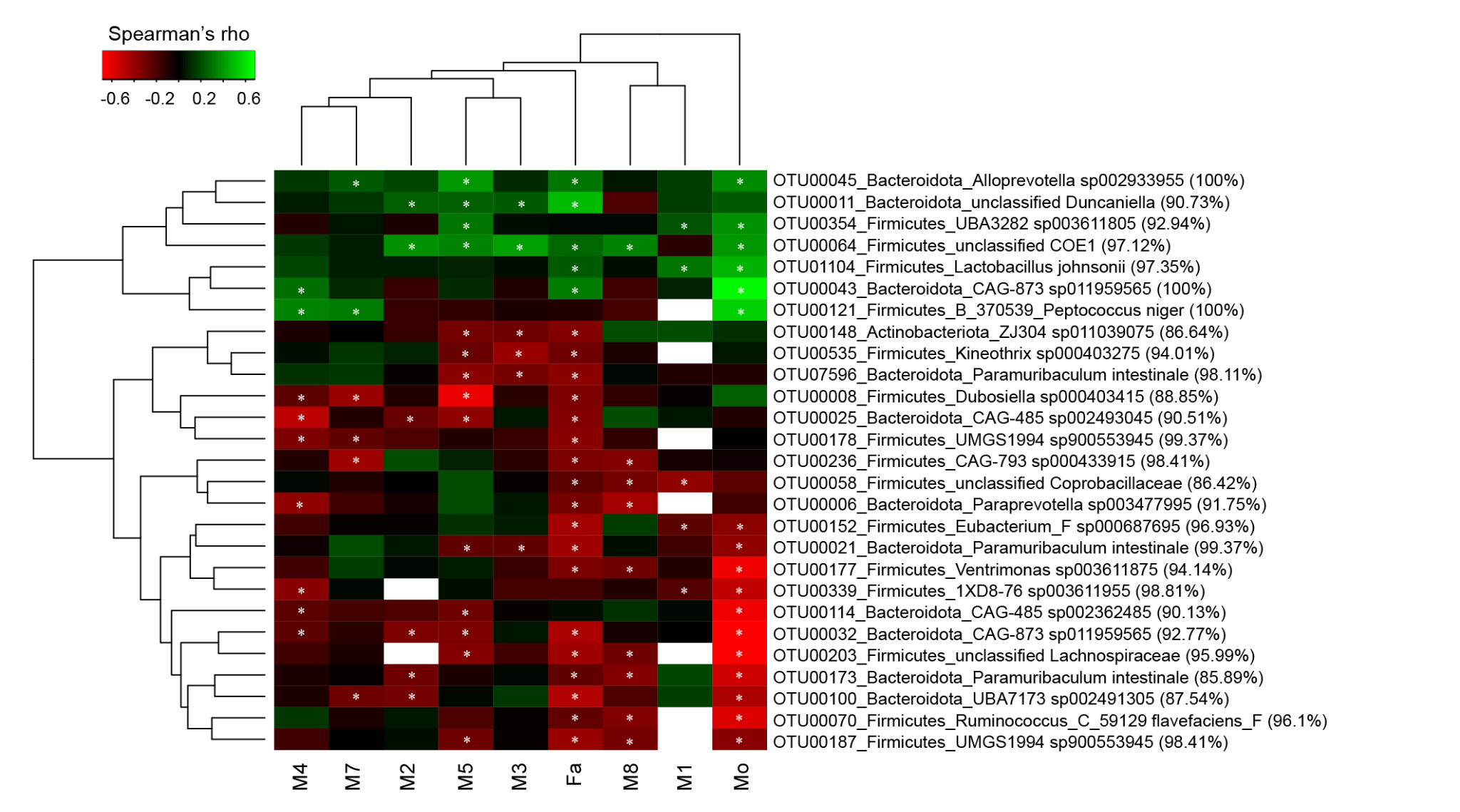

**Supplementary Fig. 13.** OTUs correlated with body weight. Heatmap of Spearman’s correlation coefficients between body weight and OTU abundance from day 555 to immediately before death (Spearman’s correlation, * *P* < 0.05).

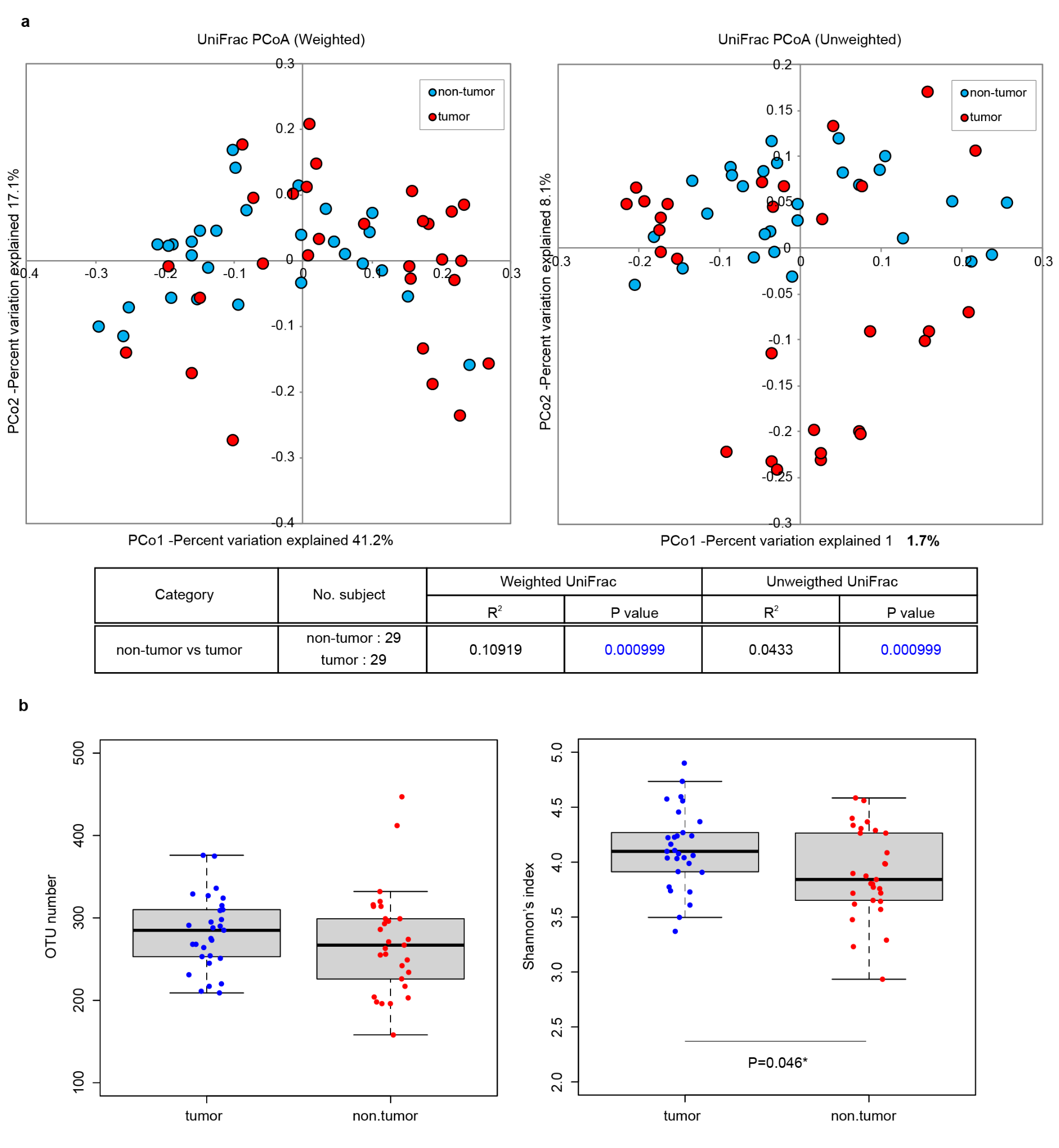

**Supplementary Fig. 14.** Effect of tumorigenesis in the last 30 days of life. **a.** Principal coordinate analysis (PCoA) of microbiome composition from 30 days before death to the last samples based on weighted/unweighted UniFrac distance. Permutational multivariate analysis of variance (PERMANOVA) results between the two groups (non-tumor, tumor) based on the post-mortem autopsy diagnoses are shown below. **b.** Boxplot of α-diversities between the tumor and non-tumor groups in the last 30 day**s** (Wilcoxon rank sum test, * *P* < 0.05)**.**

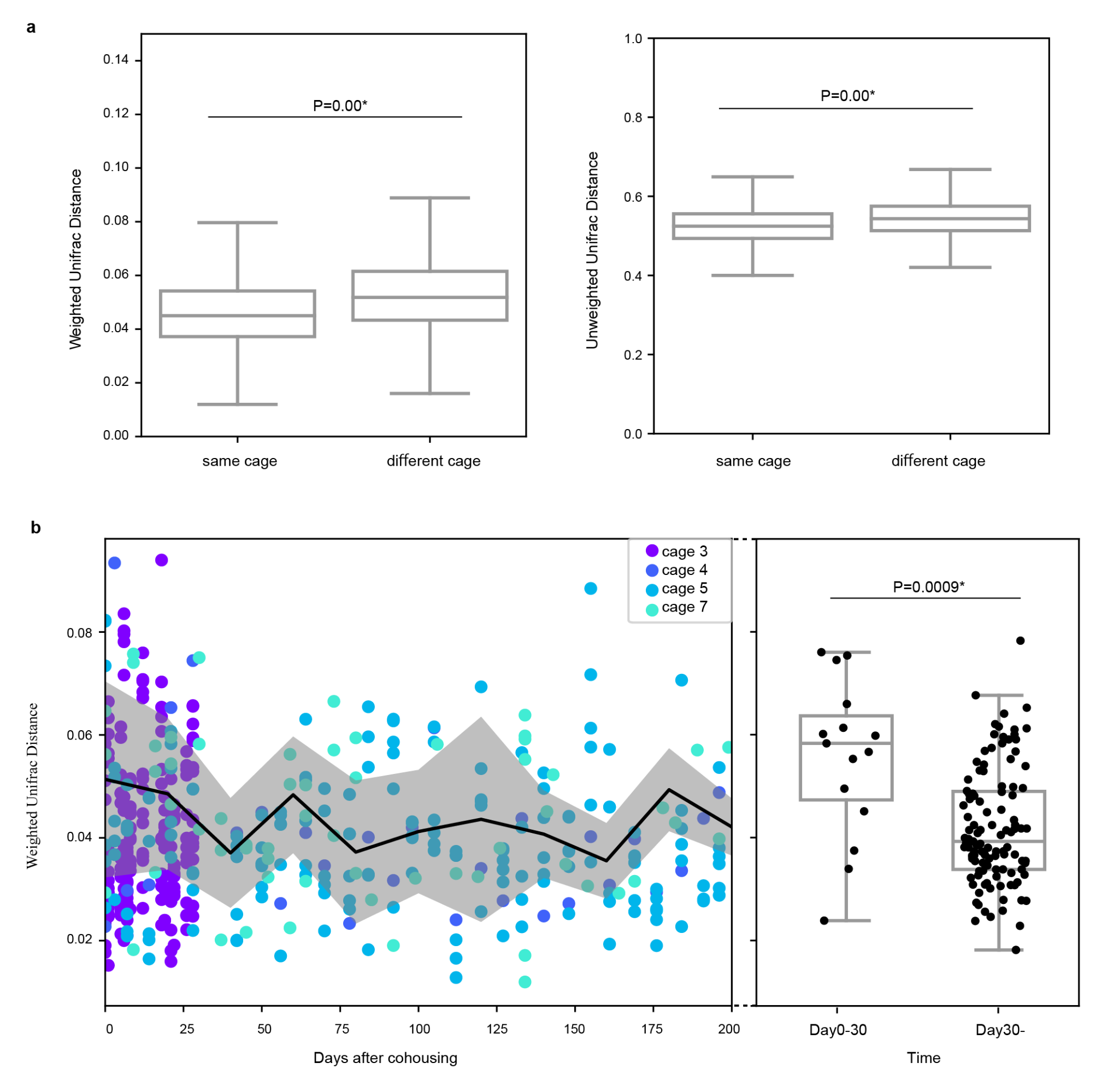
**Supplementary Fig. 15.** Cohousing cage effect. **a.** Boxplots of weighted/unweighted distances between points from the same cage and different cages are shown. The samples after day 20 were used regardless of individual differences (Wilcoxon rank sum test, * *P* < 0.05). **b.** (left) Transition of inter-individual weighted UniFrac distances inside the same cages over time. Here, day 0 is the day of starting co-housing. (right) Boxplot of the weighted UniFrac distances inside cages on days 0–30 and after day 30. The vertical axis is shared with the left figure.
